# Age is an intrinsic driver of inflammatory responses to malaria

**DOI:** 10.1101/2024.11.26.625574

**Authors:** Jessica R. Loughland, Nicholas Dooley, Zuleima Pava, Arya SheelaNair, Dean Andrew, Peta Tipping, Peter Bourke, Christian Engwerda, J. Alejandro Lopez, Kim Piera, Timothy William, Bridget E Barber, Matthew Grigg, Nicholas M Anstey, Gabriela Minigo, Michelle J. Boyle

## Abstract

Age is a critical factor influencing the host immune response to infection and disease pathogenesis. In malaria, the risk of severe disease increases with age in non-immune individuals. Malaria severity is in part driven by inflammation, but the specific cells and mechanisms contributing to age-dependent disease risk are incompletely understood. Here, we assessed inflammatory cytokines in non-immune children and adults with clinical malaria, and the phenotypic, functional and transcriptional differences of in vitro innate cell responders to malaria parasites in naive children and adults. During naturally acquired malaria, age was associated with increased plasma levels of inflammatory chemokines CCL2, CCL3, CXCL8, CXLC9, along with CRP, and IDO, which were associated with clinical symptoms. In malaria naive individuals, classical monocyte and Vδ2^+^ γδ T cell responses from adults were characterized by higher inflammatory cytokine production, and transcriptional activation following stimulation with malaria parasites. Classical monocyte responses in adults were dominated by CCL2 production, while in children the response had increased IL10 production and enrichment in IL10 signaling pathways upon parasite stimulation. This heightened inflammatory response in adults was not mitigated by parasite induced Tregs. Taken together, these findings identify cellular mechanisms of age-dependent host responses that play crucial roles in driving inflammatory responses in malaria.

## Introduction

Age is an intrinsic factor in the host response to infection, with implications for disease pathogenesis and severity (1, 2). A recent systematic analysis revealed that for many infections, disease severity follows a ‘J’ or ‘U’ curve with age, with a relatively high risk in infants followed by the lowest risk in older children, before severity risk increases again in adults and the elderly (3). The immune mechanisms underpinning these age dependent changes are largely unknown. For malaria, caused by *Plasmodium falciparum* parasite infection, the global clinical burden predominantly occurs in children, who experience the highest risk of severe disease (3). In high-transmission endemic areas, children are repeatedly infected before developing immunity which protects into adulthood, with disease severity following an ‘L’ distribution (3–5). However, in areas where malaria control has reduced transmission intensity, the burden of severe disease shifts from infants to older children (6, 7). Further, in low-transmission regions with unstable malaria, exposure and clinical immunity in childhood is much less common, and clinical disease, severe malaria and fatal *falciparum* malaria is seen in both children and adults (8). As seen for other infections, host age in these non-immune populations is an intrinsic factor in malaria pathogenesis and disease severity (9). In malaria-naive migrant populations moving from malaria-free to endemic areas, risk of severe disease is higher in adults compared to children during initial infection (10, 11). Consistent with this, amongst patients with severe disease, risk of death is higher in adults compared to children (12). Age-dependent mechanisms contributing to the risk of severe malaria are incompletely understood.

Acquisition of adaptive immunity increases with exposure, concurrently with age, complicating our ability to identify age-intrinsic mechanisms of disease risk. Indeed, few studies have compared innate immune inflammatory responses in children and adults with naturally acquired falciparum malaria, which is only possible in low transmission areas where immunity is also low, and all ages are susceptible to malaria. Clinical symptoms of malaria occur during the blood stage of parasite infection, where activation of immune cells and production of inflammatory mediators contribute to disease (13). Important innate immune cell responders include classical monocytes (14), and the innate lymphocyte γδ T cell subset expressing Vγ9Vδ2 TCR (Vδ2 T cells) (15). These cells produce key inflammatory cytokines which are associated with severe disease (16), and control of these inflammatory responses is associated with tolerogenic immunity (17). For example, in immune Malian adults there is an expansion of regulatory monocytes which produce IL10 in response to malaria (18). This expansion is malaria-driven, with Malian children having monocytes phenotypically and functionally similar to malaria-naive adults (18). However, no malaria-naive children were included in this study, so the influence of age on the monocyte response to malaria parasites is unknown. Similarly, reduced risk of malaria disease following repeat falciparum infection is associated with a reduction in the inflammatory cytokine response by γδ T cells (19, 20). However, age associated changes to Vδ2^+^ γδ T cells have also been reported, with increased inflammatory and cytotoxic potential expanding markedly after birth (21–23). The impact of these age-driven changes on the ability of malaria-naive individuals to respond to *Plasmodium* parasites is unknown.

Here, leveraging a unique cohort of children and adults with falciparum malaria from a pre-elimination low transmission area (24), we identify inflammatory cytokines which are associated with age and clinical symptoms. To dissect cells mechanisms contributing to age-dependent inflammation during malaria, we examined monocyte and Vδ2^+^ γδ T cells responses to parasite stimulation *in vitro* using cells from malaria-naive children and adults. Together, data dissected the impact of host age on the innate immune cell responses to *P. falciparum* parasites.

## Results

### Inflammatory cytokine levels correlate with age in naturally acquired malaria

Malaria disease severity, partly driven by inflammation, is associated with age in non-immune populations (6, 7). In malaria endemic areas with high transmission, immunity develops with age, and adults rarely experience symptomatic disease (4). To examine the effect of age on malaria-induced inflammation during natural infection, we analyzed inflammatory markers in plasma samples collected from children and adults who were enrolled in previously completed studies in a low malaria transmission area in Malaysia, where all ages remain susceptible to disease due to the limited prior exposure (24) (total n=97, 78 male (80%), age 21[15-45] years (median[IQR]), n=79 from patients presenting with malaria at Kudat Division district hospital (24), and n=18 enrolled at a tertiary referral hospital for the West Coast and Kudat division (25), **Supplementary Table 1)**. Amongst malaria patients, 73 (75.3%) had uncomplicated malaria, with the remainder having severe disease. Amongst malaria patients in this sample set, parasitemia was not associated with age (**Figure 1a**, *rho*=0.055, *p=*0.59). Plasma concentrations of 13 analytes related to immune inflammation were analyzed, revealing a positive correlation among inflammatory markers CCL2, CCL3, CCL4, CXCL8, CXCL9, CXCL10, as well as IDO and IL10 (**Figure 1b**). These cytokines also correlated with parasitemia, consistent with a role of parasite burden in inflammation and disease (**Figure 1b)**. Age was significantly correlated with CRP, CCL2 (MCP-1), CCL3, CXCL8, CXLC9, and IDO (**Figure 1b/c)**. Age associations were not driven by severe disease, with all correlations remaining significant when considering only patients with uncomplicated malaria (**Supplementary Figure 1a**). To assess if age-dependent inflammatory cytokines may contribute to disease, associations between disease symptoms and analytes were explored. Inflammatory analytes that were significantly associated with age, were also higher in patients with symptoms of rigors (**Figure 1d**), myalgia (muscle pain) (**Figure 1e**), headache (**Supplementary Figure 1b**) and arthralgia (joint pain) (**Supplementary Figure 1c**). CRP was significantly higher in patients with these symptoms, while chemokines CCL2 (MCP-1), CCL3 (MIP-1α) and CXCL8 (IL-8) were significantly higher in patients with rigors and myalgia (**Supplementary Figure 1b-d**). These relationships with symptoms were maintained when only analyzing patients >12 years of age, to account for potential reduced self-reporting of symptoms in children (**Supplementary Figure 1d)**. We also examined age relationships with objective clinical findings. Serum alanine aminotransferase (ALT) data was available for 62 individuals (64%) and was weakly associated with increasing age (*rho*=0.24, *p=*0.06) and CCL2 (*rho*=0.22, *p=*0.081) (**Supplementary Figure 1e/f**). Elevated ALT, a clinical measure of liver function, has been linked to parasite load and inflammatory responses during uncomplicated malaria caused by *P. falciparum* (26) and *P. vivax* (27) in populations with little pre-existing immunity. However, these findings suggest that patient age may also play a significant role in ALT elevations in non-immune individuals. Finally, we re-analyzed all clinical data from district hospital cohort, previously compared to other malaria species infections (24), for difference in clinical presentation based on age (**Supplementary Table S2).** Among symptoms at enrollment, rigors, headache, myalgia and arthralgia were more frequent in adults consistent with increased inflammation and associations between inflammatory markers and symptoms. However, as mentioned this may be confounded by higher reported symptoms in adults. The objective clinical measure of oxygen saturation was also lower in adults, consistent with increased clinical disease with age. Together data highlight the relationship between age, inflammation and disease in malaria.

**Figure 1:**
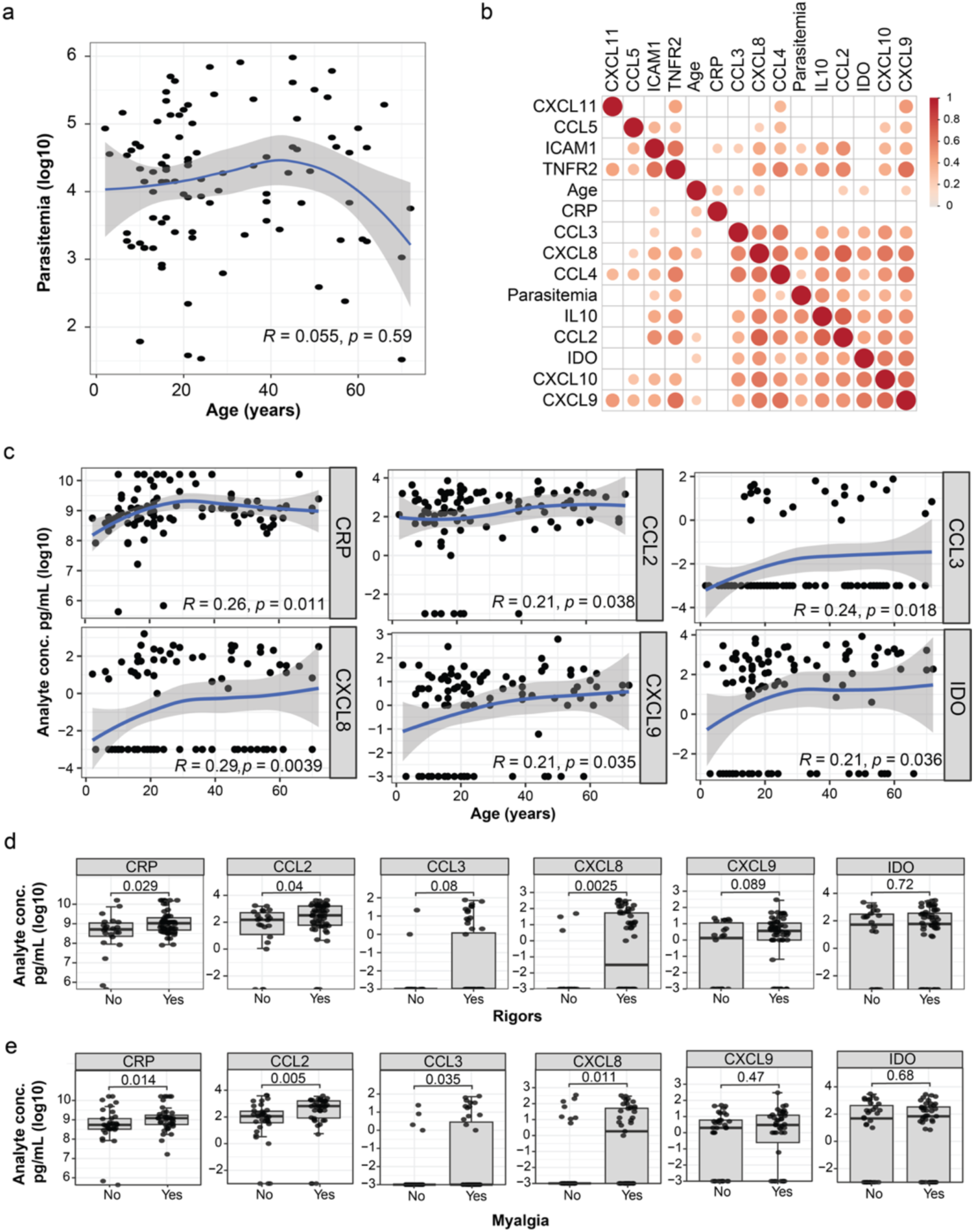
Inflammatory cytokines increase with age in patients with malaria. Thirteen analytes were measured in plasma from 97 individuals with acute malaria (78 male (80%), age 21[15-45] years (median[IQR])). **(a)** Correlation plot of patients’ parasitemia (log10 parasites/uL) and age (years). **(b)** Spearman’s correlation plot between inflammatory cytokines (pg/mL(log10)), age (years) and parasitemia (log10 parasites/μL). Correlations with p>0.05 are blank and point size scales with Rho **(c)** Individual correlation plots of analytes with significant associations with age in plot **b.** Spearman’s *Rho* and *p* are indicated. Analyte concentration for patients with and without symptoms **(d)** rigors or **(e)** myalgia (muscle pain). Tukey boxplots show the median, 25^th^ and 75^th^ percentiles of individual analytes concentration pg/mL(log10). The upper and lower hinges extend to the largest and smallest values respectively, but no further than 1.5* IQR from the hinge. Dots represent individuals. Mann-Whitney U test was used for comparisons between absence of symptoms and presence of symptoms. See also Supplementary Figure 1.

### Inflammatory cytokine production in monocytes following malaria stimulation is higher in naive adults

To understand cellular responses that may mediate increased inflammation during malaria, we analyzed phenotypes, functions and transcriptional activation profiles of immune cells relevant to malaria responsiveness in a cohort of malaria-naive individuals (children, <12 years, *n*=13, age 8 [3-12] years (median[IQR]), 38% female; adults *n*=13, age 42 [29-46] years (median[IQR]), 54% female, **Supplementary Table S3**). Analysis of *ex vivo* cell frequencies and phenotypes identified several differences, including higher proportions of CD14^+^ (classical) monocytes, CD16^+^ (non-classical) monocytes, classical dendritic cells (cDCs) and natural killer cells (NKs) in adults and a higher proportion of Vδ2^+^ γδ T cells in children (**Supplementary Figure 2**).

Among CD4^+^ T cells, adults had a higher proportion of Tfh cells, while other CD4^+^ effector T cells and FoxP3^+^ regulatory T cell (Treg) proportions were similar (**Supplementary Figure 2d-e**). Age dependent differences in expression of markers associated with activation and function (HLA-DR, CD86, CD16 and ICOS) were also detected (**Supplementary Figure 2f-h**). There was increased MHC-class II (HLA-DR) and Fc receptor (CD16) expression by classical DCs in adults but decreased co-stimulatory marker CD86 in cDCs, plasmacytoid DCs and classical monocytes (**Supplementary Figure 2d, f-h**). Activation of Vδ2^+^ γδ T cells (CD86) and Tregs (HLA-DR and ICOS) was also higher in adults compared to children (**Supplementary Figure 2d, f-g, i**).

Classical monocytes are key innate cell responders to malaria parasites (14) and are major contributors of the inflammatory cytokines associated with severe disease (16). Additionally, monocytes are major contributors of the age associated chemokines/cytokines present in the clinical data set (**Figure 1c**), including CCL2 (28), CCL3/CCL4 (29) and IL10 (30). As such, we investigated if monocytes responded to *P. falciparum* parasites in an age dependent manner. To investigate the impact of age on classical monocyte responses to malaria, PBMCs from malaria-naive children and adults were stimulated with *P. falciparum-*infected red blood cells (pRBCs) and production of IL10, IL6, IL1β, TNF and CCL2 quantified by intracellular staining (children *n=*13, adults *n=*12, **Supplementary Figure 3a**). The majority of parasite-induced cytokines were from classical monocytes, which could be distinguished from non-classical monocytes based on CD64 expression (**Supplementary Figure 3b/c**). CD64 has stable expression in culture, unlike CD16 which is down regulated (31). Stimulation with pRBCs increased cytokine production in classical monocytes compared to uninfected red blood cells (uRBCs) in both children and adults for all the cytokines tested (**Figure 2a**). However, the magnitude of parasite-induced cytokines was higher in adults for CCL2 (**Figure 2b**), while children produced significantly higher IL10 following activation (**Figure 2b**). Further, amongst all cytokine-producing monocytes, the composition of parasite-driven cytokines differed significantly between adults and children. Although both children and adults had high CCL2 production, children had a significantly more diverse cytokine response, while adults had a CCL2-dominant response (**Figure 2c**). Indeed, polyfunctionality of classical monocytes was significantly lower in adults, who had a higher proportion of monocytes producing only a single cytokine (**Figure 2c, Supplementary Figure 3d**). Furthermore, there was a positive association between age and monocytes that produced only one cytokine, and a negative association between increasing age and monocytes that produced two cytokines (**Figure 1d**). This age-associated pattern was also observed for monocytes producing three and four cytokines (**Supplementary Figure 3e**), suggesting monocyte polyfunctionality is age-dependent. While age differences in the distribution of monocyte subsets have been reported previously (32), amongst our cohort there was no difference in the proportion of classical, non-classical, or intermediate monocytes between children and adults (**Supplementary Figure 3f**). Together, these data show that classical monocyte responses to malaria parasites in naive individuals are strongly influenced by age, with adults having a focused inflammatory response while, children had a polyfunctional monocyte cytokine response with higher IL10.

**Figure 2:**
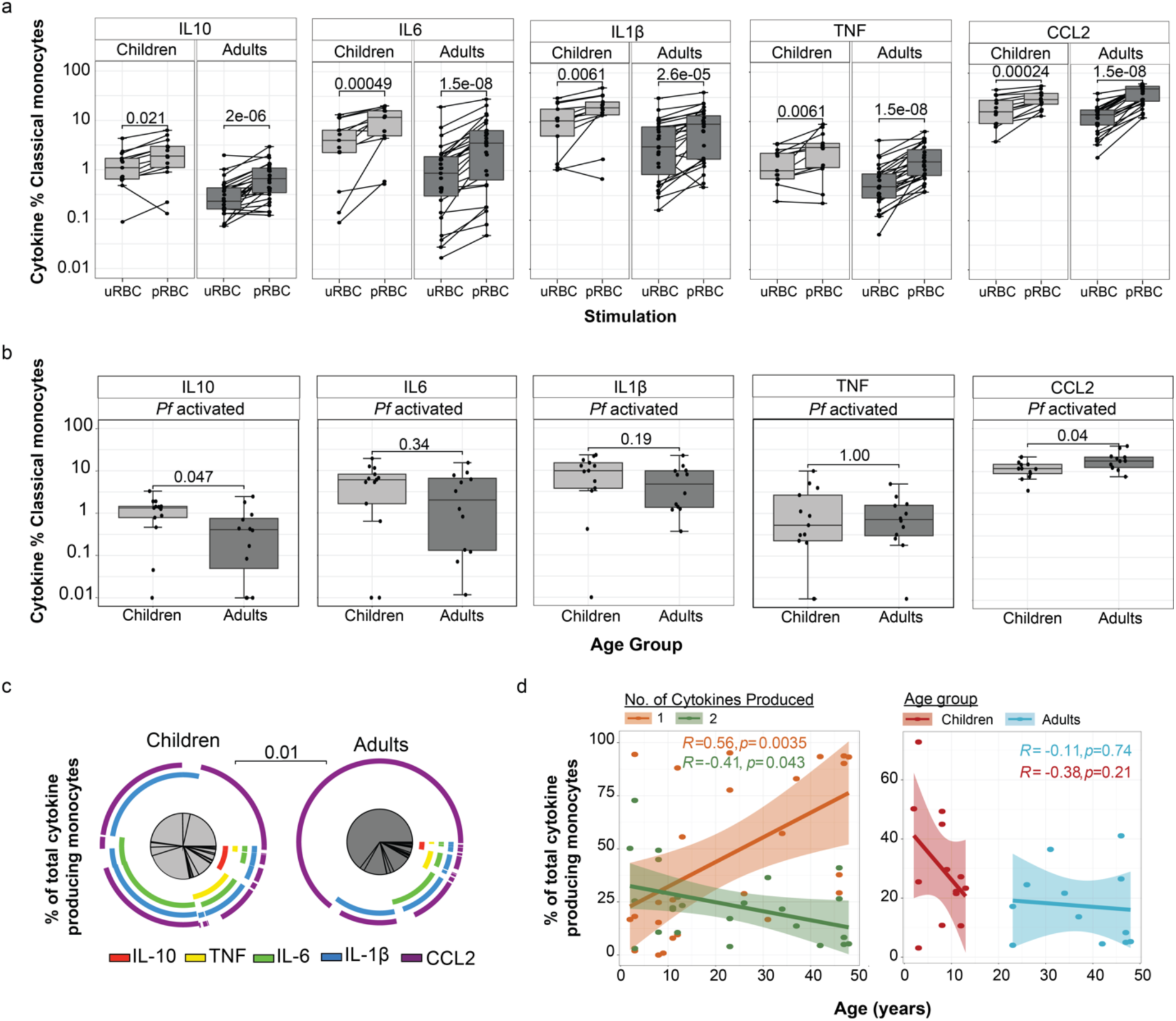
Higher inflammatory cytokine production in response to P. falciparum in monocytes from malaria naive adults compared to children. **(a)** Intracellular production of cytokines by CD14^+^ classical monocytes was analysed following stimulation with *P. falciparum-*infected (pRBCs) and uninfected (uRBCs) red blood cells with intracellular staining (children *n=*13, adults *n=*12). **(b)** Proportion of *Pf* activated cytokine expressing classical monocytes in children. Data is cytokine positive frequency in pRBC condition subtracted by uRBC condition. in each individual. **(c)** Composition of cytokines from classical monocytes following parasite stimulation. **(d)** Left plot: correlation between number of cytokines produced by monocytes and age (years). One cytokine is orange, and two cytokines are green. Right plot: correlation between monocytes which make two cytokines and age. Adults are blue, children are red. Spearman’s *Rho* and *p* are indicated. Tukey boxplots show the median, 25^th^ and 75^th^ percentiles. The upper and lower hinges extend to the largest and smallest values, respectively but no further then 1.5* IQR from the hinge. Lines represent paired observations, uRBC and pRBC comparisons are Wilcox rank paired test. Children and adult comparisons were made using the Mann-Whitney U test. Pie comparisons performed by Permutation test. Age associations were visualized by linear regressions and compared using Spearman’s rank correlation. See also supplementary figure 3.

### The transcriptional response to malaria parasites in classical monocytes is more inflammatory in naive adults

To explore the pathways driving the increased inflammatory response in monocytes from adults and increased immune regulatory response in children, we assessed the transcriptional profile of classical monocyte cells *ex vivo* and following *in vitro* stimulation with *P. falciparum* malaria parasites (**Supplementary Figure 4a/b**). Data were analysed using a general linear mixed model (glmmSeq (33)) to identify differentially expressed genes (DEGs) with age (children compared to adults), malaria stimulation (before compared to after stimulation), and genes which respond in an age-specific manner (significant for age and stimulation, and/or with a significant interaction between age and stimulation). Using this approach, a total of 10,562 DEGs were identified (**Figure 3a, Supplementary Data 1**). Malaria stimulation resulted in a large transcriptional change, with 10,395 DEGs identified. 26% of these (2,735 DEGs) were also differentially expressed with age or had a significant interaction term between age and stimulation, indicating a large age dependent response to malaria *in vitro* (**Figure 3a, Supplementary Data 1**). Principal Components Analysis (PCA) revealed differences between adults and children, as well as before and after parasite stimulation (**Figure 3b, Supplementary Figure 4c**). Age-dependent gene differences exemplified the polarization of response, with upregulated genes having a larger increase in adults, and down regulated genes having a larger decrease in children (**Figure 3c**). DEGs were categorized into 14 groups, based on whether genes were higher or lower in children or adults prior to and after stimulation, and whether expression increased or decreased in response to parasites (**Figure 3d**). The largest group of genes (876 DEGs) were higher at baseline in children but increased in adults only in response to stimulation, while decreasing in children. The second largest group (619 DEGs) were genes that were higher in children at baseline, and increased in both children and adults, but with a larger increase in adults. Both gene groups resulted in higher expression in adults after malaria parasite stimulation (**Figure 3d**). Of all DEGs which were upregulated in response to stimulation, 1,624 resulted in higher expression in adults and only 149 were higher in children (**Figure 3d**). Consistent with higher baseline (uRBC control) cytokine production in children compared to adults for all cytokines (**Figure 2a**), the majority of *Plasmodium*-responsive adult genes were higher in children *ex vivo* (2,036 vs 699 DEG).

**Figure 3:**
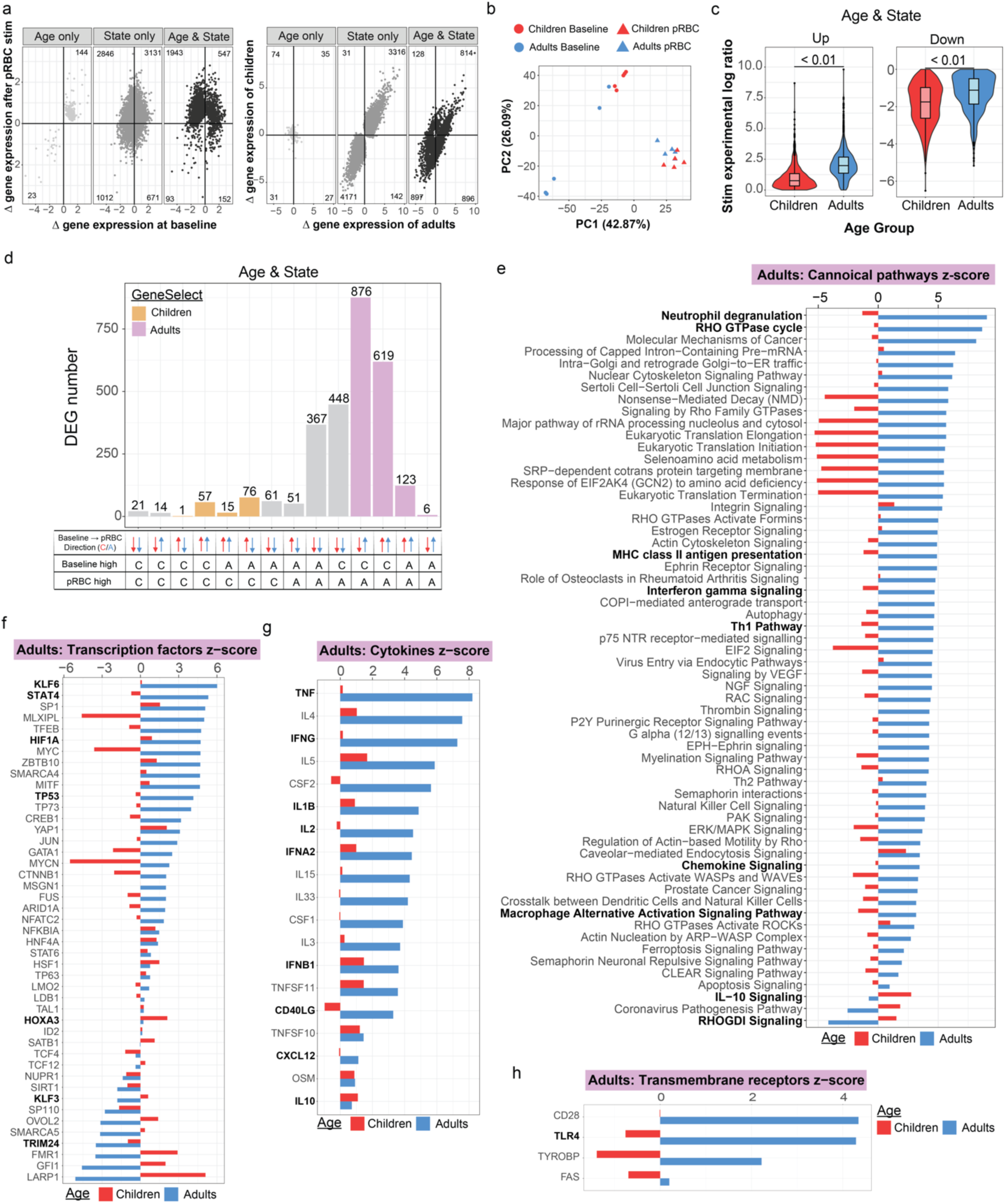
Increased transcriptional activation in monocytes from naive adults compared to children after P. falciparum parasite stimulation. (a) Scatter plots describing the shape of these transcriptional data, DEGs grouped based on if they were significant for “Age only”, “State only” or “Age & State”. Left plots, show gene expression at baseline compared to after pRBC stimulation and right plots, show gene expression levels in adults (*n=*5) compared to children (*n=*5). **(b)** Principal Components Analysis (PCA) of DEGs significant for age and state, malaria-naive volunteers before (baseline) and after pRBC stimulation (stim) **(c)** Violin plots of log ratio values of DEGs which changed with state (stimulation) and were age dependent (different between children and adults), increased (up) decreased (down) after stimulation. Comparisons by Mann-Whitney U test. **(d)** Age and State DEG counts grouped based on direction of change (up or down) and whether expression levels were higher in children or adults at baseline and/or after pRBC stimulation (post-stim). Orange bars indicate genes that increased expression after stimulation and were higher in children. Purple bars indicate genes that increased expression after stimulation and were higher in adults. Grey bars are genes that have downregulated gene expression after pRBC stimulation. **(e)** Ingenuity pathway analysis (IPA) performed using the log ratio value for children or adults and the FDR q-value of the DEGs that were upregulated following stimulation and were higher in adults (blue bars, d). Top 60 significant pathways depicted. Upstream regulator analysis showing predicted **(f)** transcription factors, **(g)** cytokines and **(h)** transmembrane receptors to be activated or inhibited in adults and children, using DEGs as in e. Benjamin-Hochberg corrected P-values used to identify significant pathways and upstream regulators in the IPA analysis. See also supplementary figures 4/5.

We performed Ingenuity Pathway Analysis (IPA) of all upregulated DEGs that were higher in adults following pRBC stimulation. Most enriched pathways were uniquely enriched in adults (**Figure 3e**). Consistent with increased inflammatory cytokine responses in adults, adult-specific upregulated pathways included inflammatory IFNγ signaling pathways, MHC Class II presentation, as well as chemokine signaling and the macrophage alternative activation signaling pathway (**Figure 3e**). Furthermore, consistent with significantly higher IL10 production by monocytes in children, the IL10 signaling pathway was upregulated in children. Further, Rho GDP-dissociation inhibitor (RHOGDI) signaling was also inhibited in adults but activated in children. This pathway refers to the regulatory mechanisms involved in modulating the activity of Rho GTPases which regulate inflammatory responses (34). The top upregulated genes in adults after stimulation included the chemokine *CXCL11* and *SLAMF9* (**Supplementary Figure 4d**). *CXCL11,* which was also higher *ex vivo* in adults (**Supplementary Figure 4d**), has been shown to increase in circulation during malaria, particularly during a primary infection (35) and has roles in malaria pathogenesis (29). *SLAMF9,* which was higher in adults after parasite stimulation, promotes the initiation of an inflammatory response in antigen-presenting cells such as monocytes (36). Additionally, adults had higher expression of the MHC Class II gene *HLA-DRA* after stimulation, consistent with a more robust response with higher functional potential (**Supplementary Figure 4d**). Consistent with IPA, upstream regulators predicted to be uniquely activated in adults were also indicative of an increased inflammatory response to malaria parasites (**Figure 3f**). Transcriptional factors increased in adults compared to children included KLF6, which drives pro-inflammatory gene expression (37), and STAT4 which is a critical driver of inflammation (38). In contrast in children, upregulated transcriptional factors included HOXA3 which inhibits M1 polarization and drives M2 polarization (39), and KLF3, a suppressor of monocyte inflammation (40) (**Figure 3f).** Inflammatory cytokines were predicted to be more activated in adults, including IFNγ, TNF, IL6, IL1β and type 1 IFNs (IFNα and IFNβ). In addition, CD40LG, IL-2 and CXCL12 were predicted to be activated in adults and inhibited in children. IL10 was predicted to be activated in both children and adults but was higher in children (**Figure 3g**). Further, within the transmembrane receptor category of upstream regulators, Toll-Like Receptors 4 (TLR4), a pathogen recognition receptor that can bind to GPI anchors derived from the malaria parasite (41), was activated in adults and inhibited in children (**Figure 3h)**, consistent with an increased ability of monocytes from adults to bind parasite products and induce inflammation. In contrast, of the 149 DEGs (5%) which were higher in children after parasites stimulation (**Figure 3d**), only two pathways were detected by IPA, with minimal differences in children and adults in upstream regulators (**Supplementary Figure 5a-d**). Taken together, transcriptional and cytokine production data show that classical monocytes’ responsiveness to malaria is age dependent, with enhanced inflammatory responses in adults compared to children in malaria naive individuals. This finding is consistent with the major role of monocytes in malaria pathogenesis (14) and the increased risk of severe disease in adults during primary infection (11).

### Inflammatory cytokine production from Vδ2^+^ γδ T cells following malaria stimulation is higher in naive adults

Inflammatory transcriptional signatures in monocytes included the chemokine CXCL11, which has been linked to expansion of Vδ2^+^ γδ T cells in adults during a primary *P. falciparum* infection (35). Vδ2^+^ γδ T cell, along with monocytes, are major producers of inflammatory cytokines associated with disease severity in malaria (16). Following stimulation with *P. falciparum* parasites *in vitro*, Vδ2^+^ γδ T cells from both children and adults responded robustly to parasite stimulation with increased production of IFNγ and TNF (**Figure 4a, Supplementary Figure 6a/b**). There was a significantly higher production of both IFNγ and TNF in adults (**Figure 4b**). Cytokines production was from both single-producing and IFNγ^+^/TNF^+^ co-producing Vδ2^+^ γδ T cells (**Figure 4b**/c). Within children, the frequency of IFNγ and TNF following stimulation significantly increased between 0-10 years, highlighting the age dependent increase in inflammatory Vδ2^+^ γδ T cells (**Figure 4d**).

**Figure 4:**
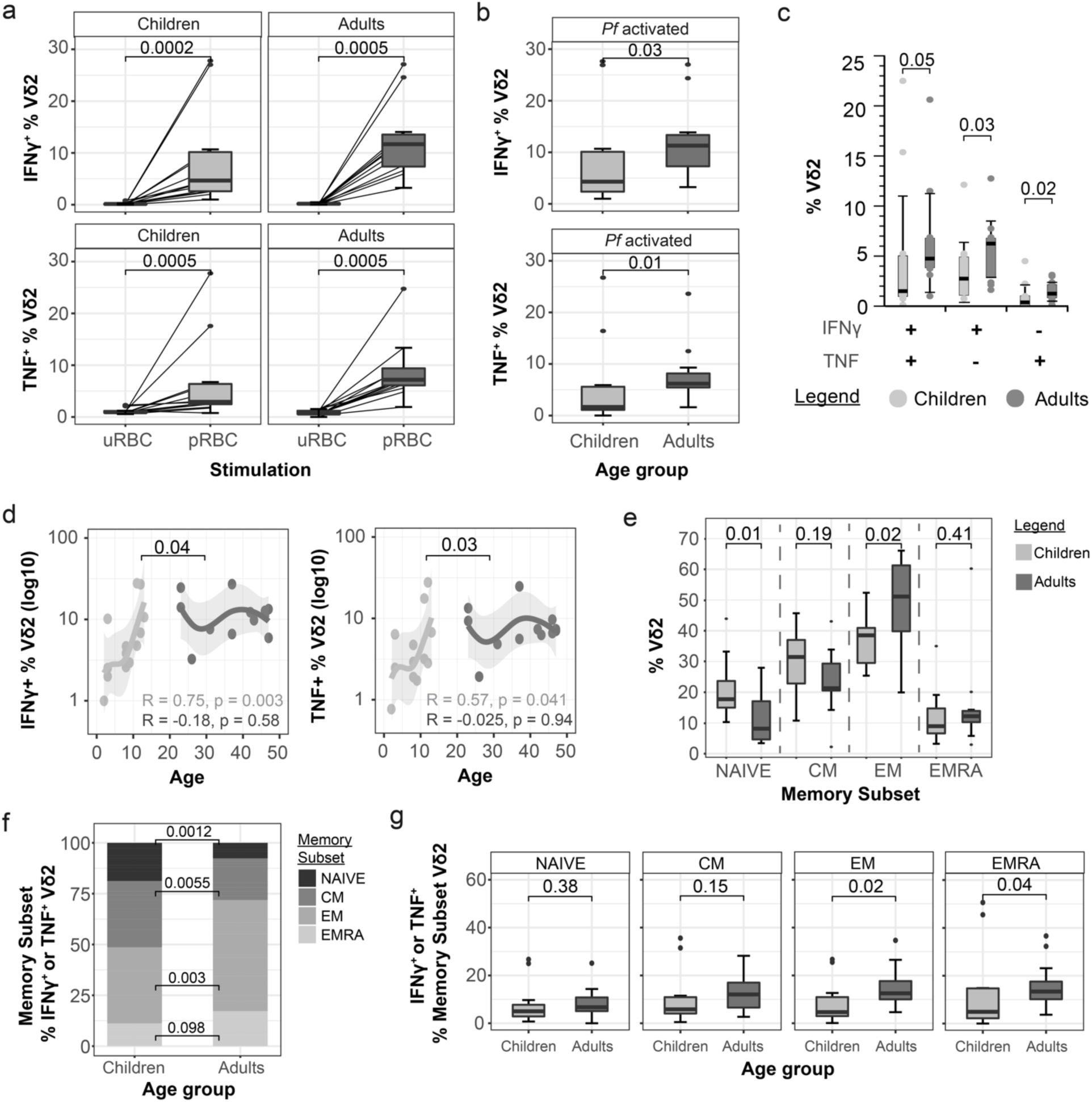
Malaria-naive adult Vδ2^+^ γδ T cells produce more inflammatory cytokines following malaria stimulation. **(a)** Intracellular production of IFNγ and TNF by Vδ2^+^ γδ T cells was analysed following 24-hour stimulation with *P. falciparum* infected (pRBCs) and uninfected (uRBCs) red blood cells with intracellular staining (children *n=*13, adults *n=*12). **(b)** Comparison of children *versus* adult parasite specific cytokine response. Data is cytokine positive frequency in pRBC condition subtracted by uRBC condition in each individual. **(c)** Co-expression of IFNγ and TNF as a percentage of total Vδ2^+^ γδ T cells (parasite specific response). **(d)** IFNγ and TNF positive Vδ2^+^ γδ T cell frequency to pRBC stimulation was positively associated with age in years and separated into groups of children and adults. **(e)** Vδ2^+^ γδ T cell memory subset frequency comparisons after pRBC stimulation between children and adults: NAIVE (CD27^+^CD45RA^+^), CM (CD27^+^CD45RA^-^), EM (CD27^-^CD45RA^-^) and EMRA (CD27^-^CD45RA^+^). **(f)** Memory subset proportions of IFNγ or TNF producing Vδ2^+^ γδ T cells in children and adults. **(g)** Comparisons of total *Pf* activated cytokine production of cells producing either IFNγ or TNF within each Vδ2^+^ γδ T cell memory subsets between children and adults. Tukey boxplots show the median, 25^th^ and 75^th^ percentiles. The upper and lower hinges extend to the largest and smallest values, respectively but no further than 1.5* IQR from the hinge. Lines represent paired observations. Wilcoxon signed rank test was used to match paired data. Children and adult comparisons were made using the Mann-Whitney U test. Age associations were visualised by loess regressions and compared using Spearman’s rank correlation. See also Supplementary Figure 6.

The memory subset distribution of Vδ2^+^ γδ T cells has been reported to change with age, with a reduction in naive (CD27^+^ CD45RA^+^) Vδ2^+^ γδ T cells with age (22, 23). Consistent with this, the proportion of effector memory Vδ2^+^ γδ T cells (EM; CD27^-^ CD45RA^-^) was significantly higher in adults compared to children (**Figure 4e**). EM Vδ2^+^ γδ T cells were the dominant malaria responding subset, producing more IFNγ and TNF compared to other cell subsets (naive [NAIVE]; central memory [CM]; CD27^+^ CD45RA^-^; and terminally differentiated effector memory [EMRA]; CD27^-^ CD45RA^+^) following stimulation (**Figure 4f**). This suggests that the increased response in adults may be due to age-dependent changes to Vδ2^+^ γδ T subset distribution. However, when IFNγ and TNF production within each memory subsets was examined, both EM and EMRA Vδ2^+^ γδ T cells produced significantly more cytokines in adults compared to children (**Figure 4g**). As such, the increased cytokine response of Vδ2^+^ γδ T cells in adults is due to both the expansion of responsive effector memory cells within the Vδ2^+^ γδ T cell compartment, and an increased capacity of responding cells to produce cytokine.

### Transcriptional response to malaria parasites in Vδ2^+^ γδ T cells is similar in naive children and adults

As for monocytes, we next examined the transcriptional profile of purified Vδ2^+^ γδ T cells *ex vivo* and following stimulation with parasites to identify pathways mediating increased inflammation in adults (**Figure 5a, Supplementary Data 2**). As seen for monocytes, Vδ2^+^ γδ T cells had a large transcriptional response to parasite stimulation, with 8,766 DEGs identified after stimulation, of which 35% (3,062) were also significant for age, or had a significant interaction with age (**Figure 5a**). However, in contrast to classical monocytes, Principal Components Analysis revealed fewer differences between children and adults both before and after stimulation in the Vδ2^+^ γδ T cells (**Figure 5b, Supplementary** Figure 7a). Furthermore, DEGs upregulated with stimulation had larger increases in children, and down regulated genes had larger decreases in adults (**Figure 5c**). When DEGs were grouped based on expression before/after stimulation, and directional change, only 710 genes were increased and were higher in adults following stimulation compared to 2,015 DEGs which increased with stimulation and were higher in children (**Figure 5d**). However, when these two gene sets were analysed by IPA, largely similar pathways with comparable enrichment scores were identified (**Figure 5e, Supplementary Figure 7b/c**). Of pathways with higher enrichment in children, the majority of these were involved in translation and metabolism. (**Figure 5f**). Similar upstream regulators were predicted to be enriched in Vδ2^+^ γδ T cells between adults and children (**Supplementary Figure 7d-f)**.

**Figure 5:**
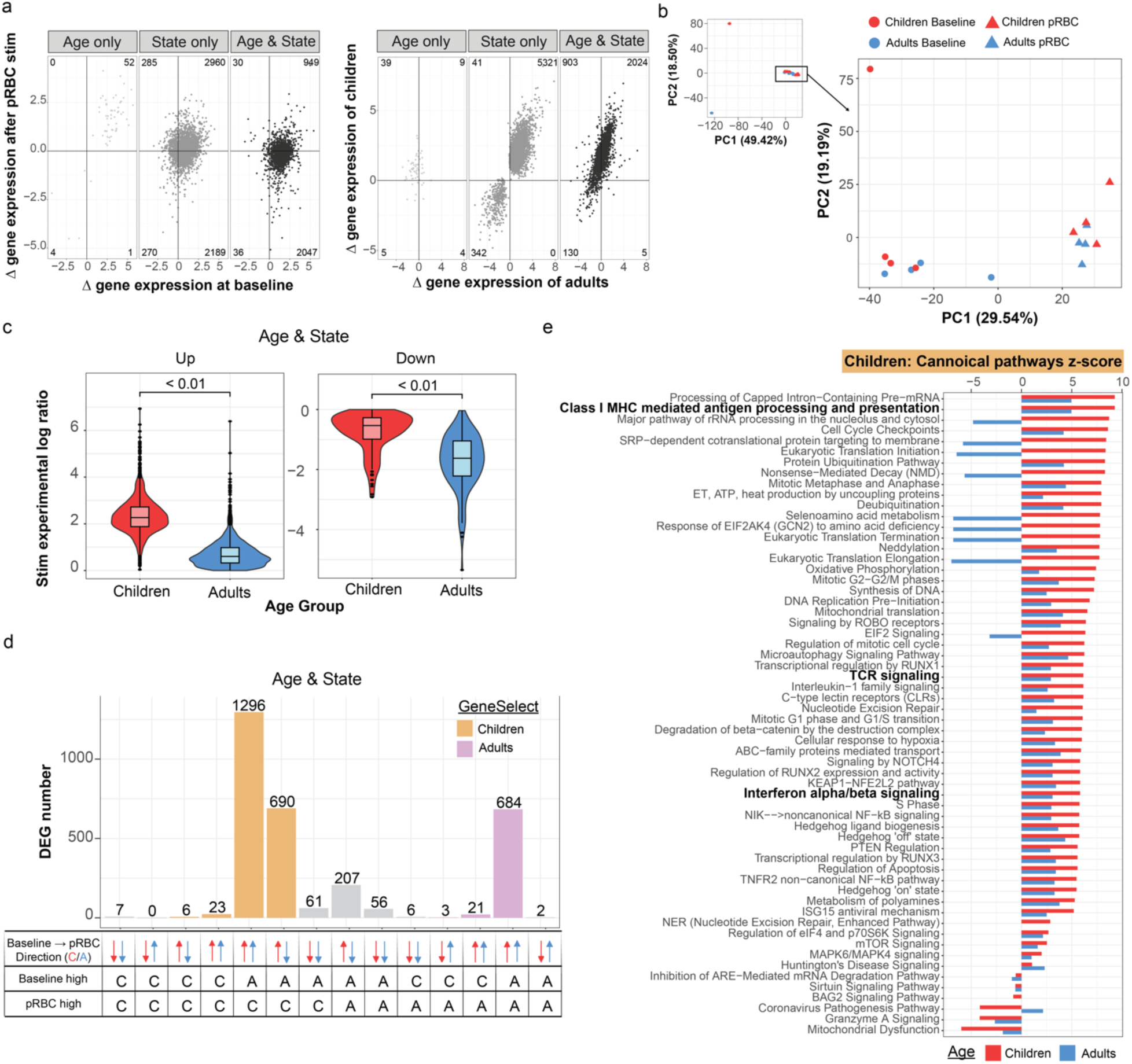
Transcriptional activation of Vδ2^+^ γδ T cells by malaria is similar in naive children and adults. **(a)** Scatter plots describing the shape of these transcriptional data, DEGs grouped based on if they were significant for “Age only”, “State only” or “Age & State”. Left plots, show gene expression at baseline compared to after pRBC stimulation and right plots, show gene expression levels in adults (*n=*5) compared to children (*n=*5). **(b)** Principal Components Analysis (PCA) of DEGs significant for age and state, malaria-naive volunteers before (unstim) and after parasite stimulation (stim) **(c)** The log ratio values of the genes that went “up” after pRBC stimulation and “down” after pRBC stimulation. **(d)** Barplot grouping the DEGs based on their direction of change (up or down) and whether expression levels were higher in children or adults at baseline and after pRBC stimulation. Purple bars indicate genes that increased expression levels after stimulation and were higher in adults compared to children after stimulation. Orange bars indicate genes that increased expression levels after stimulation and were higher in children compared to adults after stimulation. Grey bars are genes that have downregulated gene expression after pRBC stimulation. **(e)** Ingenuity pathway analysis (IPA) performed using the log ratio value for children or adults and the FDR q-value of the DEGs that were upregulated following stimulation and were higher in children (red bars, d). Top 60 significant pathways depicted. Benjamin-Hochberg corrected P-values used to identify significant pathways and upstream regulators in the IPA analysis. See also supplementary figure 7.

Interestingly, gene expression levels of IFNγ and TNF were significantly increased after parasite stimulation but were not different between age groups (**Supplementary Data 2**). The disconnect between transcriptional and functional data may be due to the differences in timing of assays, or due to post-translational age-dependent control of inflammation and IFNγ/TNF expression.

### Increased innate cell inflammatory responses in naive adults is not balanced by induction of FOXP3^+^ Tregs

Inflammation from innate immune cells in response to pathogens requires control to avoid immunopathogenesis. In malaria, control of inflammation can be mediated by multiple cell responses, including Tregs (42). Tregs expand during primary *Plasmodium* infection in adults, and expansion is associated with reduced inflammation (43). Activation of Tregs has been linked to the immunoregulatory enzyme indoleamine 2,-dioxygenase (IDO) (44, 45). In our data IDO was age associated in clinical malaria in plasma (**Figure 1b-c)** and was significantly upregulated in adults following parasite stimulation in the RNAseq monocyte data set **(Supplementary Figure 4d)**, suggesting that adults may induce higher Treg cell frequencies to control increased inflammation. Tregs can be expanded from malaria naive individuals following parasite stimulation *in vitro* (46–50). As such, we tested whether age impacted the malaria induced expansion of Tregs in naive individuals, hypothesizing that increased inflammation in response to malaria parasites in adults would be balanced by increased induction of Tregs. Following stimulation, Tregs were identified as CD4^+^/CD25high/CD127low/Foxp3^+^ T cells, activation measured by CD38, ICOS or Ki67 and inhibition potential measured by CCR4, TNFR2 and PD-1 expression (51) (**Figure 6a, Supplementary Figure 8a/b**). Treg expansion occurred to comparable levels in both children and adults (**Figure 6b**). Parasite expanded Treg populations had increased expression of activation markers compared to control-stimulated cells, but with no significant difference between age groups (**Figure 6c**). Further, while the expression of inhibitory markers was higher in adults in cells stimulated with either parasite infected or uninfected RBCs (**Supplementary Figure 8c**), there was no difference in the magnitude of induction of inhibitory markers between adults and children (**Figure 6d**). Within these data, we also investigated whether the inflammatory CD4 T cell subset (Th1) was expanded or activated during in vitro pRBC stimulation (**Supplementary Figure 9a/b**). IFNγ production from CD4 T cells is associated with parasite clearance, but also disease (52). Similar to Tregs, Th1 (CXCR3^+^ CCRR6-) CD4T cells had increased expression of activation markers (**Supplementary Figure 9c**) and inhibition markers compared to control-stimulated cells (**Supplementary Figure 9d**).

**Figure 6:**
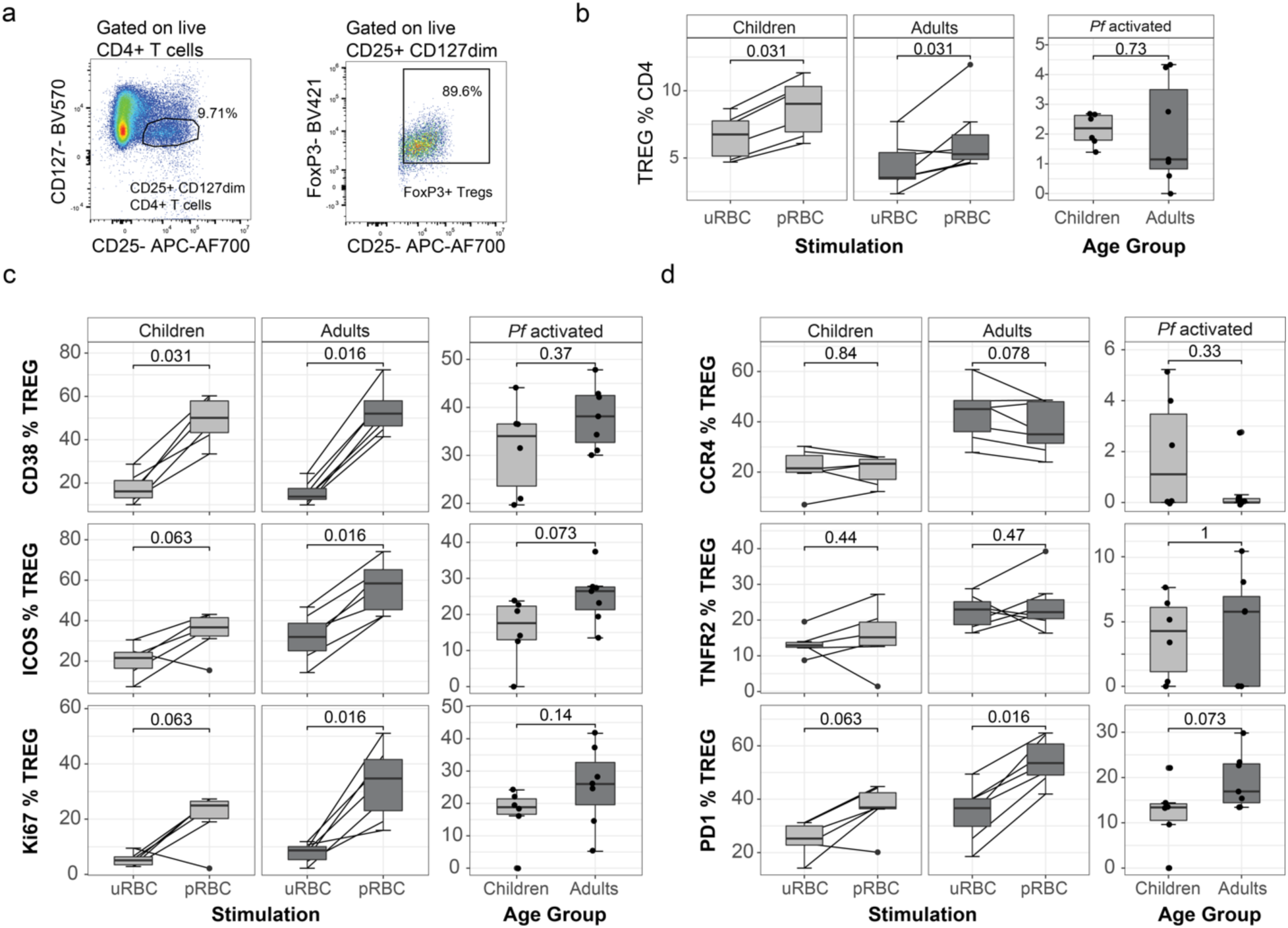
Comparable Foxp3^+^ Treg response in both adults and children in response to malaria. CD4^+^ T regulatory cell (Treg) expansion and surface marker expression in children (*n=*6) and adults (*n=*7) was measured after 5 days *in vitro* co-culture with trophozoite stage *P. falciparum* red blood cells (pRBCs) and uninfected (uRBCs) red blood cells. (**a**) Tregs were identified as CD25high CD127low and FoxP3^+^ CD4 T cells. **(b)** Treg frequency within CD4^+^ T cells following stimulation. Surface marker frequency of inflammatory immune activation (**c**: CD38, ICOS and Ki67) and inhibition (**d**: CCR4, TNFR2 and PD-1) post stimulation of Treg frequency with uRBC, pRBC and *Pf* activated responses (quantified as expression in pRBC minus expression in uRBC cultured conditions). Tukey boxplots show the median, 25^th^ and 75^th^ percentiles. The upper and lower hinges extend to the largest and smallest values, respectively but no further than 1.5* IQR from the hinge. Lines represent paired observations. Wilcoxon signed rank test was used to compare paired data. Children and adult comparisons were made using the Mann-Whitney U test. See also supplementary Figures 8/9.

However, responses were comparable between children and adults, suggesting that the increased inflammatory responses in monocytes and γδ T cells is not a global enhancement of inflammation with all cells of the immune response but rather specific to innate cells.

## Discussion

Here we identify an age-dependent inflammatory response to *P. falciparum* both during human disease and *in vitro* that may contribute to age-dependence of malaria pathogenesis and disease severity risk (9, 11, 12). In a cohort of patients with clinical malaria, plasma levels of inflammatory cytokines were associated with age and disease symptoms. This is despite age being independent from circulating parasitemia. In unexposed individuals, both monocytes and Vδ2^+^ γδ T cells produced significantly higher inflammatory cytokines in adults compared to children following *in vitro* parasite stimulation. Higher inflammation was not balanced by more robust induction of Treg cells, nor was there age associated differences in Th1 CD4 T cell responses. Together with previous studies in the same clinical population showing age-dependence of endothelial activation in falciparum malaria (9), our findings suggest that age-dependent immune mediated inflammation contributes to age-dependent disease risk in malaria.

In the absence of migration of naive individuals into areas of disease risk (11), or emergence of new pathogens such as SARS-CoV-2, understanding age intrinsic factors influencing the immune response to infection is confounded by prior exposure and the development of adaptive immunity. Thus, while it is well recognized that the immune system evolves with age, with important consequences for disease risk (1–3), few studies have defined age specific mechanisms between children and adults in responses to individual pathogens. Here, we identify that both monocytes and Vδ2^+^ γδ T cells are significantly more inflammatory in adults compared to children in response to malaria. These cells have been shown to be major producers of cytokines associated with severe malaria (16). Monocytes produce inflammatory cytokines to malaria parasites via several pathways (14), including via TLR2 and TLR4 recognition of Glycosylphosphatidylinositol (GPI) anchors (41), TLR8 responding to *Plasmodium* RNA (53), and via TLR4 binding hemozoin bound with parasite proteins (54). Previous studies have shown that monocyte responsiveness to TLR2 and TLR4 stimulation varies with age, but responsiveness to TLR7/8 is age-independent (55, 56). This suggests that the age dependent response to malaria may be driven by specific pattern recognition pathways. Indeed, data presented here suggests that increased inflammatory responses to malaria parasites may be mediated by increased TLR4 activation, which was transcriptionally activated in monocytes in adults and not children in response to parasite stimulation.

Amongst top upregulated genes in monocytes from adults following parasite stimulation was the chemokine CXCL11, which acts as a chemotactic for activated T cells and is increased in symptomatic malaria compared to asymptomatic infected individuals (57). CXCL11, along with CXCL9 has also been linked to expansion of Vδ2^+^ γδ T cells in adults during a primary *P. falciparum* infection (35). Consistent with this link, Vδ2^+^ γδ T cells from malaria naive adults have a higher inflammatory response to malaria following *in vitro* stimulation compared to children and circulating CXCL9 was correlated with age in patients with malaria. This increased inflammatory response was both due to the expansion of EM cells within the Vδ2^+^ γδ T cell population, and increased reactivity of all memory subsets to parasite stimulation. Along with roles of Vδ2^+^ γδ T cells during infection, these cells also have been associated with protection induced by whole parasite irradiated sporozoite vaccines (58, 59). While these vaccines induce sterile protection in malaria-naive adults, they have failed to induce any efficacy in infants in endemic areas with this failure hypothesized to be linked to Vδ2^+^ γδ T cell function (60). Our data supports this hypothesis by identifying a clear age dependent responsiveness of γδ T cells to malaria parasites.

In contrast to responses in adults, cell responses in children had increased regulatory potential, particularly in monocytes, with both transcriptional enrichment of regulatory pathways including IL10 and RHODGI signaling in children, as well as increased production of IL10 in response to parasite stimulation. In malaria endemic areas, responses from monocytes isolated from immune Malian adults produce more IL10 and less inflammatory cytokines (TNF, IL6 and IL-1b) compared to both non-immune children, and malaria unexposed adults (18), consistent with the induction of immunoregulatory networks by malaria exposure associated with protection from disease (17). Here we show that in naive populations, monocytes from children also produce more IL10 compared to adults, in an age intrinsic manner, distinct from malaria induced regulator networks. In contrast, while frequencies and functions of Tregs cells are highest early in life (61), we found no evidence that induction of Tregs by malaria parasites was age dependent. However, it should be noted that circulating IDO levels were increased with age in patients with malaria, so increased Treg induction may still occur during clinical disease. Further studies are required to investigate the age dependent induction of other regulatory cell responses during malaria, such as Tr1 CD4 T cells which co-produce IL10 with IFNγ and dominate the CD4 T cell compartment during malaria infection (62, 63).

While increased inflammatory response with age may increase disease severity and risk, this heightened responsiveness may also be benefitable for the development of adaptive immunity. In malaria naive children and adults moving into high endemic areas, while adults had initially increased risk of severe disease, they acquired protection from infection more rapidly (11, 64).

This age-dependent acquisition of anti-parasitic immunity, largely mediated via adaptive responses, has also been shown in Ugandan children in high transmission areas (65). We have previously linked increased antibody development after malaria in adults to increased activation of T-follicular helper cells (66). Further studies are required to investigate innate and adaptive responses during malaria across age.

Limitations of our study include the absence of both young infants and the elderly in both our unexposed healthy and natural infection cohorts. Rapid maturation of the immune system in infants may further modulate the age responsiveness to malaria reported here and would likely show differences to the children studied here. Indeed, changes to TLR stimulation in myeloid cells rapidly adapt in the first year of life (67), and γδ T cells gain effector functions in the months after birth (21). Further, risk of severe disease also increases in older adults, compared to young adults (9, 12, 68), thus future studies are required to understand innate cell responsiveness to malaria parasites in these age groups. Additionally, no cellular samples were available from our clinical cohort of natural infection, thus we were not able to investigate age-dependent changes to monocytes and γδ T cell activation during clinical malaria. Further, due to limited cell numbers in malaria naive donors, we were also unable to investigate other innate cell responses which may have roles in malaria disease risk.

In conclusion, these data identify classical monocytes and Vδ2^+^ γδ T cells as important drivers of the inflammatory response in malaria-naive adults upon first exposure to malaria parasites. Age-dependent inflammatory responses and similar age-dependence in endothelial activation (9), may contribute to the increased risk of severe disease in adults compared to children in low malaria transmission settings. These findings have important implications for our understanding of the distinct immune response in children and adults upon first exposure to a pathogen. In addition to informing age dependence of inflammation in malaria, our findings may be of relevance to other infectious diseases were severity follows ‘J’ or ‘U’ shape age distributions (2, 3)

## Materials and Methods

### Ethics statement

Written informed consent was obtained from all study participants or, in the case of children, parents or guardians. Studies were approved by the ethics committees of the Northern Territory Department of Health and Menzies School of Health Research (Darwin, Australia, HREC 2010-1431, HREC-2012-1766), Medical Research and Ethics Committee, Ministry of Health, Malaysia (NMRR-10-754-6684 and NMRR-12-499-1203), QIMR-Berghofer Human Research Ethics Committee (HREC P3445 and P3444) and the Alfred Hospital Ethics Committee (HREC 188/23 and 80/24).

## Study participants

### Clinical cohorts

Age related circulating cytokines/chemokines and clinical associations were analyzed in all available participants from patients (n=76) enrolled previously published studies of patients reporting with *P. falciparum* malaria at district hospitals in Kudat Division, northwest Sabah, Malaysia (24), and in an additional 18 individuals enrolled at a referral hospital for the West Coast and Kudat Division (25) (**Supplementary Table S1**). Patients presenting to study hospitals with microscopy-diagnosed malaria of any species were enrolled following written informed consent. Children were predefined as age <12 years, based on Malaysian Ministry of Health pediatric ward admission guidelines. For the current analysis, patients were included if *Plasmodium* species PCR-confirmed *P. falciparum* mono-infection. Laboratory analysis for clinical parameters was conducted as previously described in parent studies. Clinical data from the district hospital cohort (24) was reanalyzed based on age with children defined as <12 years, compared to adults >12 years (**Supplementary Table S2**). Here, plasma samples collected from lithium heparin blood collection tubes during enrollment were used.

#### Malaria-naive children and adults

Blood samples were collected from children (*n* = 13, 61% male, age 8 [3-12] years (median [IQR])) and adults (*n* = 13, 46% male, age 42 [29-46] years (median [IQR])) from an outpatient allergen clinic at the Royal Darwin Hospital (**Supplementary Table S3**). Participants had not experienced an allergic reaction for >6 months, had not taken immune medication, were malaria-naive and were confirmed healthy by an on-site immunologist. Peripheral blood mononuclear cells (PBMCs) were separated by ficoll-paque density gradient centrifugation and cryopreserved. As expected, children had a higher absolute lymphocyte count (Children: 2.8 [2.5-3.7] x10^9^/uL (median [IQR]); Adults: 2.1 [1.7-2.5] x10^9^/uL (median [IQR]), *p=*0.002), while there was no difference in the whole blood monocyte count (Children: 0.5 [0.5-0.6] x10^9^/uL (median [IQR]); Adults: 0.5 [0.4-0.6] x10^9^/uL (median [IQR])).

### Luminex Assay

ProcartaPlex^TM^ human custom kits were purchased from Thermo Fisher Scientific (MA, USA) and run according to manufacturer’s recommendations with the following modifications.

Cryopreserved plasma from individuals with naturally acquired malaria during acute infection were diluted 3X before performing assay with a custom kit (Cat #: PPX-27-MXNKUV6) and 30 000X for human simplex CRP kits (Cat #: EPX01A-102 88-901). After sample dilution, 25 uL of plasma was incubated with 25uL of 1X UAB ((Cat #: EPX-11111-000) and magnetic capture beads at 4°C overnight on an orbital shaker set to 600rpm. Plates were washed twice then incubated with Biotinylated detection antibody at RT for 30 mins at 600 rpm. Plates were washed twice then incubated with Streptavidin-PE at RT for 30 mins at 600 rpm. Plates were washed twice, resuspended in Reading Buffer then acquired on the Bio-Rad Bio-Plex 200 array reader. Wells with a bead count of ≥50 were included in analysis. Optimised standard curve 4PL/5PL regressions were calculated with Bio-Plex Manager software and used to convert bead Fluorescence intensity to concentration (pg/mL) dependent on the standard concentrations provided by the manufacturers. Sample analyte values beyond the ULOQ or below the LLOQ were set to ULOQ ^+^ 1 and LLOQ –1, respectively.

### Parasite Culture

Packed red blood cells (RBCs) from donors were infected *in vitro* with the *Plasmodium falciparum* 3D7 parasite strain (Trager, 1976). Packed RBCs for parasite culture were acquired from the Australian Red Cross. *Plasmodium falciparum* infected RBCs (pRBCs) were cultured at 5% haematocrit in Roswell Park Memorial Institute 1640 media (RPMI) supplemented with AlbuMAX II (0.25%) and heat-inactivated human sera (5%). Cultures were incubated at 37 °C in 1% O2, 5% CO2, 94% N2 gas mixture. Culture media was replaced daily, and parasite stage/parasitemia was monitored by Giemsa-stained blood smears. pRBCs were grown to 15% parasitemia and purified from uninfected RBCs (uRBCs) and early stage pRBCs, via magnet separation to enrich mature trophozoite stage pRBCs. Purified pRBCs (>95% purity) were stored at -80 °C following addition of a Glycerolyte cryopreservant.

### Flow Cytometry

#### Ex vivo PBMC phenotyping

*Ex vivo* PBMC phenotype and activation were assessed by flow cytometry. PBMCs were thawed in 10% FBS/RPMI, and surface staining performed in 2% FBS/PBS using fluorescent-tagged antibodies to identify cell lineages and measure activation marker expression of interest (**Supplementary** Figure 2 **a/b, Supplementary Table S4)**.

#### Innate cell pRBC stimulation assay

PBMCs were co-cultured at 37 °C, 5% CO2 in a 96-well U-bottom plate for 24 hours at a 1:1 pRBC:PBMC ratio with 1x10^6^ mature trophozoite stage pRBCs or 1x10^6^ uRBCs, prepared as previously described. Protein transport inhibitors (Monensin, BD GolgiStop) were added after 1 hour at 37 °C, 5% CO2. At 24h, cells were stained to identify monocytes and Vδ2 T cells (**Supplementary Table S5**), washed with 2% FCS/PBS, cells were permeabilised with 1 X Perm/Wash^TM^ (BD Biosciences) and stained for intracellular cytokine production (**Supplementary Table S5**). To determine the pRBC specific response uRBC responses were subtracted from pRBC responses.

#### Treg/ CD4 T cell expansion assay

To measure *in vitro* Treg activation and expansion we performed a 5 day PBMC co-culture with pRBC at a 3:1 cell ratio. After 5 days, cells were washed with 2% FCS/PBS and surface stained to identify Tregs and Treg activation (**Supplementary Table S6**). To determine the pRBC specific response uRBC responses were subtracted from pRBC responses. FACS data was acquired using the Cytek Aurora 3 laser (CA, USA), and analysed with FlowJo v10 (BD, 2019).

### Cell isolation and RNA sequencing

PBMCs were thawed in 10% FBS/RPMI. Classical monocytes and Vδ2 T cells subsets were FACS sorted using the BD FACSAria™ III Cell Sorter from *ex vivo* PBMC or after *in vitro* co-culture at 37 °C, 5% CO2 in 96-well U-bottom plates for 4 hours at a 1:1 cell ratio with 1x10^6^ mature trophozoite stage pRBCs. RNA was extracted from isolated cell population lysates using the QIAGEN PicoPureTM RNA isolation kit (Applied Biosystems™, KIT0204), and RNA quality confirmed with the 2200 TapeStation system (G2964AA) by High Sensitivity RNA ScreenTape (5067–5579). RNA sequencing libraries were constructed using the NEBNext® Single Cell/Low Input RNA Library Prep Kit for Illumina® (E6420S) and NEBNext® Multiplex Oligos for Illumina® (96 Unique Dual Index Primer Pairs) (E6440S). The libraries were sequenced using a paired-end NextSeq 500/550 high output kit v2.5 (150 cycles) (Cat number 20024907).

## Data Analysis

### Flow cytometry

For the *ex vivo* phenotyping panel we used the R package SPECTRE (69) to identify cell subsets of interest. Unsupervised clustering was performed using both children (*n=*12) and adult (*n=*11) PBMC sample, and cell clusters were visualised with uniform manifold approximation and projection (UMAP) (**Supplementary Figure 2a/b, Supplementary Table S2**). Expression of lineage markers were used to annotate cell clusters into 12 high level cell states (**Supplementary Figure 2a/b**). For functional assays, data was analysed with FlowJo v10 (BD, 2019) and R/R studio was used for statistical analysis and data visualisation.

### Bulk-RNAseq

Raw sequencing reads were first trimmed to remove adapter sequences and low-quality bases using Cutadapt (v1.9). Trimmed reads were then aligned to the Human GRCh37 reference genome, incorporating Ensembl v97 gene models, using STAR (v2.5.2a). Alignment files were processed, sorted, and converted to the required formats using SAMtools (v1.9). Gene and transcript expression levels were quantified with RSEM (v1.2.30), providing normalized expression estimates. Quality assessment of the RNA-seq data was performed using RNA-SeQC (v1.1.8) to ensure data reliability. All analyses were carried out using Python (v3.6.1) and Perl (v5.22) for scripting and workflow automation. Low-count genes (fewer than 10 counts) were removed, and dispersion was estimated using edgeR (v4.40) workflow in R. To compare gene expression in paired pre- and post-stimulated samples from children and adults, we used negative binomial mixed-effects models with the glmmSeq R package (33). The design matrix accounted for random effects of individual samples (**Supplementary** Figure 4b) and included an interaction term between state (pre- vs. post-stimulation) and age (adult vs. child). Fold change due to stimulation was calculated by subtracting the response term of unstimulated from stimulated samples. Fold change due to age was calculated by subtracting the response term of children from adults for both stimulated and unstimulated samples. We corrected for multiple testing using Storey’s q-value method, defining significance for q-values lower than 0.05 with the q-value R package.

Ingenuity Pathway Analysis (IPA) (Qiagen, Hilden, Germany) was used for canonical pathway enrichment and predicted upstream regulator analysis using DEGs with a false detection rate (FDR) < 0.05, with no fold-change cut-off.

#### Statistical analysis

Non-parametric testing was performed for all analysis. Continuous data for all cellular responses and plasma cytokines were compared between the children and adults’ groups using Mann-Whitney U test or correlated with age as a continuous variable with Spearman’s correlations. Statistical comparisons were not adjusted for multiple comparisons. All analyses were performed in R (version 4.4.4). Graphical outputs were made in ggplot2 (version 3.5.1) and ggpubr (version 0.6.0). No sample size calculation was performed, instead all available participant data was included. No subgroup analysis was performed. Data generation was performed with blinding to participant demographic data.

## Acknowledgements

We thank all participants in this study; the clinical and laboratory research staff and the hospital directors at the study sites, the head of Clinical Research Centre Malaysia and colleagues and staff of the UK medical research council *P. knowlesi* Monkeybar Project. RBC used for parasite culture were provided by Australian Red Cross Blood Bank (Brisbane). We thank the Director-General of Health, Malaysia, for permission to publish this article.

## Author contributions

Conceptualization and methodology: JRL, NLD, GM, MJB

Investigation and validation: NLD, JRL, DA, AS, BEB, MG, MJB

Formal analysis: NLD, JRL, ZP, MG, MJB

Resources: PT, PB, KP, GM, TW, BEB, MG, NMA,

Data Curation: NLD, JRL, ZP, MG, MJB.

Writing - Original Draft: NLD, JRL, MG, MJB

Supervision: JAL, CE, NMA, TW, MG, BEB, MJB

Writing - Review & Editing: NLD, JRL, GM, MJB with critical feed back and approval from all authors

All authors have read and approved the final version of the manuscript.

## Data availability

The fastq files and the raw counts were deposit in the Gene Expression Omnibus (GEO) database under the accession number GSE270553 at: https://www.ncbi.nlm.nih.gov/geo/query/acc.cgi?acc=GSE270553.

FCS files are available at http://flowrepository.org/id/FR-FCM-Z8F2

## References

1. Simon AK, Hollander GA, McMichael A. Evolution of the immune system in humans from infancy to old age. Proceedings Biological sciences. 2015;282(1821):20143085.

2. Glynn JR, Moss PAH. Systematic analysis of infectious disease outcomes by age shows lowest severity in school-age children. Sci Data. 2020;7(1):329.

3. Abel L, Casanova J-L. Human determinants of age-dependent patterns of death from infection. Immunity. 2024;57(7):1457–1465.

4. Doolan DL, Dobaño C, Baird JK. Acquired Immunity to Malaria. Clin Microbiol Rev. 2009;22(1):13–36.

5. Geneva: World Health Organisation. World malaria report 2023. 2023.

6. Griffin JT, Ferguson NM, Ghani AC. Estimates of the changing age-burden of Plasmodium falciparum malaria disease in sub-Saharan Africa. Nat Commun. 2014;5(1):3136.

7. Ghani AC, et al. Loss of Population Levels of Immunity to Malaria as a Result of Exposure-Reducing Interventions: Consequences for Interpretation of Disease Trends. PLoS ONE. 2009;4(2):e4383.

8. Poespoprodjo JR, et al. Malaria. Lancet. 2023;402(10419):2328–2345.

9. Barber BE, et al. Effects of Aging on Parasite Biomass, Inflammation, Endothelial Activation, Microvascular Dysfunction and Disease Severity in Plasmodium knowlesi and Plasmodium falciparum Malaria. J Infect Dis. 2017;215(12):1908–1917.

10. Baird JK. Age dependent characteristics of protection v. susceptibility to Plasmodium falciparum. Ann Trop Med Parasitol. 2017;92(4):367–390.

11. Baird JK, et al. Age-Dependent Susceptibility to Severe Disease with Primary Exposure to Plasmodium falciparum. J Infect Dis. 1998;178(2):592–595.

12. Dondorp AM, et al. The Relationship between Age and the Manifestations of and Mortality Associated with Severe Malaria. Clin Infect Dis. 2008;47(2):151–157.

13. Schofield L, Grau GE. Immunological processes in malaria pathogenesis. Nat Rev Immunol. 2005;5(9):722–735.

14. Dobbs KR, Crabtree JN, Dent AE. Innate immunity to malaria—The role of monocytes. Immunological reviews. 2019;293(1):8–24.

15. Dantzler KW, Jagannathan P. γδ T Cells in Antimalarial Immunity: New Insights Into Their Diverse Functions in Protection and Tolerance. Front Immunol. 2018;9:2445.

16. Stanisic DI, et al. γδ T cells and CD14+ Monocytes Are Predominant Cellular Sources of Cytokines and Chemokines Associated With Severe Malaria. J Infect Dis. 2014;210(2):295–305.

17. Boyle MJ, Engwerda CR, Jagannathan P. The impact of Plasmodium-driven immunoregulatory networks on immunity to malaria. Nat Rev Immunol. 2024;1–17.

18. Guha R, et al. Plasmodium falciparum malaria drives epigenetic reprogramming of human monocytes toward a regulatory phenotype. Plos Pathog. 2021;17(4):e1009430.

19. Jagannathan P, et al. Loss and dysfunction of Vδ2+ γδ T cells are associated with clinical tolerance to malaria. Sci Transl Med. 2014;6(251):251ra117-251ra117.

20. Jagannathan P, et al. Vδ2+ T cell response to malaria correlates with protection from infection but is attenuated with repeated exposure. Scientific Reports. 2017;7(1):11487.

21. Papadopoulou M, et al. Fetal public Vγ9Vδ2 T cells expand and gain potent cytotoxic functions early after birth. Proc Natl Acad Sci. 2020;117(31):18638–18648.

22. De Rosa SC, et al. Ontogeny of γδ T Cells in Humans. J Immunol. 2004;172(3):1637–1645.

23. Gray JI, et al. Human γδ T cells in diverse tissues exhibit site-specific maturation dynamics across the life span. Sci Immunol. 2024;9(96):eadn3954.

24. Grigg MJ, et al. Age-Related Clinical Spectrum of Plasmodium knowlesi Malaria and Predictors of Severity. Clinical Infectious Diseases. 2018;67(3):350–359.

25. Barber BE, et al. A prospective comparative study of knowlesi, falciparum, and vivax malaria in Sabah, Malaysia: high proportion with severe disease from Plasmodium knowlesi and Plasmodium vivax but no mortality with early referral and artesunate therapy. Clinical Infectious Diseases. 2013;56(3):383–397.

26. Reuling IJ, et al. Liver Injury in Uncomplicated Malaria is an Overlooked Phenomenon: An Observational Study. EBioMedicine. 2018;36:131–139.

27. Odedra A, et al. Liver Function Test Abnormalities in Experimental and Clinical Plasmodium vivax Infection. Am J Trop Med Hyg. 2020;103(5):1910–1917.

28. Gschwandtner M, Derler R, Midwood KS. More Than Just Attractive: How CCL2 Influences Myeloid Cell Behavior Beyond Chemotaxis. Front Immunol. 2019;10:2759.

29. Ioannidis LJ, Nie CQ, Hansen DS. The role of chemokines in severe malaria: more than meets the eye. Parasitology. 2014;141(5):602–613.

30. Loughland JR, et al. Transcriptional profiling and immunophenotyping show sustained activation of blood monocytes in subpatent Plasmodium falciparuminfection. Clinical & Translational Immunology. 2020;9(6):126–18.

31. Ong S-M, et al. A Novel, Five-Marker Alternative to CD16–CD14 Gating to Identify the Three Human Monocyte Subsets. Front Immunol. 2019;10:1761.

32. Neeland MR, et al. Children and Adults in a Household Cohort Study Have Robust Longitudinal Immune Responses Following SARS-CoV-2 Infection or Exposure. Front Immunol. 2021;12:741639.

33. Lewis M, Goldmann K, Sciacca E. glmmSeq: General Linear Mixed Models for Gene-Level Differential Expression. 2022.

34. Tong L, Tergaonkar V. Rho protein GTPases and their interactions with NFκB: crossroads of inflammation and matrix biology. Biosci Rep. 2014;34(3):e00115.

35. Lautenbach MJ, et al. Systems analysis shows a role of cytophilic antibodies in shaping innate tolerance to malaria. Cell Reports. 2022;39(3):110709.

36. Wilson TJ, et al. Signalling lymphocyte activation molecule family member 9 is found on select subsets of antigen-presenting cells and promotes resistance to Salmonella infection. Immunology. 2020;159(4):393–403.

37. Date D, et al. Kruppel-like Transcription Factor 6 Regulates Inflammatory Macrophage Polarization*. J Biol Chem. 2014;289(15):10318–10329.

38. Kaplan MH. STAT4: a critical regulator of inflammation in vivo. Immunol Res. 2005;31(3):231–241.

39. Al Sadoun H, et al. Enforced Expression of Hoxa3 Inhibits Classical and Promotes Alternative Activation of Macrophages In Vitro and In Vivo. J Immunol. 2016;197(3):872–884.

40. Knights AJ, et al. Krüppel-like factor 3 (KLF3) suppresses NF-κB–driven inflammation in mice. J Biol Chem. 2020;295(18):6080–6091.

41. Krishnegowda G, et al. Induction of Proinflammatory Responses in Macrophages by the Glycosylphosphatidylinositols of Plasmodium falciparum CELL SIGNALING RECEPTORS, GLYCOSYLPHOSPHATIDYLINOSITOL (GPI) STRUCTURAL REQUIREMENT, AND REGULATION OF GPI ACTIVITY*. J Biol Chem. 2005;280(9):8606–8616.

42. Scholzen A, Minigo G, Plebanski M. Heroes or villains? T regulatory cells in malaria infection. Trends Parasitol. 2010;26(1):16–25.

43. Walther M, et al. Upregulation of TGF-β, FOXP3, and CD4+CD25+ Regulatory T Cells Correlates with More Rapid Parasite Growth in Human Malaria Infection. Immunity. 2005;23(3):287–296.

44. Woodberry T, et al. Early Immune Regulatory Changes in a Primary Controlled Human Plasmodium vivax Infection: CD1c+ Myeloid Dendritic Cell Maturation Arrest, Induction of the Kynurenine Pathway, and Regulatory T Cell Activation. Infect Immun. 2017;85(6):10.1128/iai.00986-16.

45. Munn DH, Mellor AL. Indoleamine 2,3 dioxygenase and metabolic control of immune responses. Trends Immunol. 2013;34(3):137–143.

46. Scholzen A, Cooke BM, Plebanski M. Plasmodium falciparum induces Foxp3hi CD4 T cells independent of surface PfEMP1 expression via small soluble parasite components. Front Microbiol. 2014;5:200.

47. Minigo G, et al. Parasite-dependent expansion of TNF receptor II-positive regulatory T cells with enhanced suppressive activity in adults with severe malaria. PLoS pathogens. 2009;5(4):e1000402.

48. Scholzen A, et al. Plasmodium falciparum–Mediated Induction of Human CD25hiFoxp3hi CD4 T Cells Is Independent of Direct TCR Stimulation and Requires IL-2, IL10 and TGFβ. PLoS pathogens. 2009;5(8):e1000543–20.

49. Finney OC, et al. Freeze-thaw lysates of Plasmodium falciparum-infected red blood cells induce differentiation of functionally competent regulatory T cells from memory T cells. Eur J Immunol. 2012;42(7):1767–1777.

50. Finney OC, Riley EM, Walther M. Phenotypic analysis of human peripheral blood regulatory T cells (CD4+FOXP3+CD127lo/–) ex vivo and after in vitro restimulation with malaria antigens. Eur J Immunol. 2010;40(1):47–60.

51. Sakaguchi S, et al. Regulatory T Cells and Human Disease. Annu Rev Immunol. 2020;38(1):1–26.

52. Kumar R, et al. The regulation of CD4+ T cells during malaria. Immunological reviews. 2020;293(1):70–87.

53. Coch C, et al. Human TLR8 Senses RNA From Plasmodium falciparum-Infected Red Blood Cells Which Is Uniquely Required for the IFN-γ Response in NK Cells. Front Immunol. 2019;10:371.

54. Barrera V, et al. Host fibrinogen stably bound to hemozoin rapidly activates monocytes via TLR-4 and CD11b/CD18-integrin: a new paradigm of hemozoin action. Blood. 2011;117(21):5674–5682.

55. Liao S-L, et al. Maturation of Toll-like receptor 1–4 responsiveness during early life. Early Hum Dev. 2013;89(7):473–478.

56. Levy O, et al. Selective Impairment of TLR-Mediated Innate Immunity in Human Newborns: Neonatal Blood Plasma Reduces Monocyte TNF-α Induction by Bacterial Lipopeptides, Lipopolysaccharide, and Imiquimod, but Preserves the Response to R-848. J Immunol. 2004;173(7):4627–4634.

57. Che JN, et al. Chemokines responses to Plasmodium falciparum malaria and co-infections among rural Cameroonians. Parasitol Int. 2015;64(2):139–144.

58. Mordmüller B, et al. Sterile protection against human malaria by chemoattenuated PfSPZ vaccine. Nature. 2017;542(7642):445–449.

59. Zaidi I, et al. γδ T Cells Are Required for the Induction of Sterile Immunity during Irradiated Sporozoite Vaccinations. The Journal of Immunology. 2017;199(11):3781–3788.

60. Oneko M, et al. Safety, immunogenicity and efficacy of PfSPZ Vaccine against malaria in infants in western Kenya: a double-blind, randomized, placebo-controlled phase 2 trial. Nat Med. 2021;27(9):1636–1645.

61. Thome JJC, et al. Early-life compartmentalization of human T cell differentiation and regulatory function in mucosal and lymphoid tissues. Nat Med. 2016;22(1):72–77.

62. Boyle MJ, et al. Effector Phenotype of Plasmodium falciparum-Specific CD4+ T Cells Is Influenced by Both Age and Transmission Intensity in Naturally Exposed Populations. The Journal of infectious diseases. 2015;212(3):416–425.

63. Boyle MJ, et al. The Development of Plasmodium falciparum-Specific IL10 CD4 T Cells and Protection from Malaria in Children in an Area of High Malaria Transmission. Frontiers in immunology. 2017;8:1329.

64. Baird JK, et al. Age-dependent acquired protection against Plasmodium falciparum in people having two years exposure to hyperendemic malaria. The American journal of tropical medicine and hygiene. 1991;45(1):65–76.

65. Rodriguez-Barraquer I, et al. Quantification of anti-parasite and anti-disease immunity to malaria as a function of age and exposure. Elife. 2018;7:e35832.

66. Oyong DamianA, et al. Adults with Plasmodium falciparum malaria have higher magnitude and quality of circulating T-follicular helper cells compared to children. Ebiomedicine. 2022;75:103784.

67. Nguyen M, et al. Acquisition of Adult-Like TLR4 and TLR9 Responses during the First Year of Life. Plos One. 2010;5(4):e10407.

68. Schwartz E, et al. Age as a Risk Factor for Severe Plasmodium falciparum Malaria in Nonimmune Patients. Clin Infect Dis. 2001;33(10):1774–1777.

69. Ashhurst TM, et al. Integration, exploration, and analysis of high-dimensional single-cell cytometry data using Spectre. Cytom Part A. [published online ahead of print: 2021]. 10.1002/cyto.a.24350.

