## Supplementary Figures and Tables for "Age is an intrinsic driver of inflammatory responses to malaria"

**Supplementary Tables and Figures**

**Supplementary Table S1: Demographics of patients analyzed for plasma cytokines**

| **Demographics** | **Total cohort** | **District hospital**[#] | **Referral hospital**[^] |
|---|---|---|---|
| **Number** | 97 | 79 | 18 |
| **Age,** years, median [IQR], range | 21, [15-45], 2-72 | 21 [14-40.5], 2-72 | 24.5 [18.25-45.75], 14-66 |
| **Male sex**, n (%) | 78 (80%) | 61 (77%) | 17 (94%) |
| **Parasites/ µL,** median (IQR) | 18 525 (2 522-63 612) | 9 924 (2030-32 839) | 233 409 (142 012-425 315) |
| **Severe malaria**[$]**,** n (%) | 24 (24.7%) | 6 (7.6%) | 18 (100%) |

### Samples from District Hospital enrollments #ref24, Grigg MJ, et al. Age-Related Clinical Spectrum of Plasmodium knowlesi Malaria and Predictors of Severity. *Clinical Infectious Diseases*. 2018;67(3):350–359. All available samples from this cohort were tested

^ Samples from referral hospital #ref25. Barber BE, et al. A prospective comparative study of knowlesi, falciparum, and vivax malaria in Sabah, Malaysia: high proportion with severe disease from Plasmodium knowlesi and Plasmodium vivax but no mortality with early referral and artesunate therapy. *Clinical Infectious Diseases*. 2013;56(3):383–397. A subset of participants were tested selected based on plasma volume availability.

[$] Severe malaria was defined using World Health Organization 2014 research criteria.

#### Age is an intrinsic driver of inflammatory responses to malaria

**Supplementary Table S2: Malaria clinical cohort characteristics district hospital cohort**

| **Patient characteristic** | ***Children*** *≤12 yrs* | ***Adolescents/adults*** *>12 yrs* | **P-value** |
|---|---|---|---|
| **Number** (% total) | 31 (32.3) | 65 (67.7) | - |
| Age, years | | | |
| Median (IQR) | 7 (3-10) | 24 (16-47) | - |
| Range | 1-12 | 1-12 | |
| Male gender, n (%) | 21 (67.7) | 48 (73.8) | 0.534 |
| Previous malaria (self-reported), n (%) | 3 (9.7) | 8 (12.7) | 0.705 |
| Symptoms on enrolment, n (%) | | | |
| Rigors | 14 (45.2) | 49 (76.6) | **0.004** |
| Headache | 21 (67.7) | 56 (86.2) | **0.034** |
| Vomiting | 11 (35.5) | 28 (43.1) | 0.479 |
| Abdominal pain | 9 (29.0) | 18 (27.7) | 0.891 |
| Diarrhoea | 4 (12.9) | 7 (10.8) | 0.759 |
| Cough | 12 (38.7) | 22 (33.8) | 0.712 |
| Shortness of breath | 4 (12.9) | 10 (15.4) | 0.741 |
| Myalgia | 7 (22.6) | 35 (53.8) | **0.004** |
| Arthralgia | 7 (22.6) | 34 (52.3) | **0.006** |
| Examination findings on enrolment | | | |
| Oxygen saturation, %, median (IQR) | 100 (99-100) | 99 (98-100) | **0.010** |
| Parasite count, parasites/µL, median (IQR) | 7392 (1462-36546) | 9924 (2522-22860) | 0.934 |

#### Age is an intrinsic driver of inflammatory responses to malaria

**Supplementary Table S3: Malaria naive cohort information**

| | **Children <12 years** | **Adults >12 years** | **P** |
|---|---|---|---|
| Total, n (%) | 13, (50%) | 13, (50%) | |
| Sex, male, n (%) | 8, (61%) | 6, (46%) | ns[#] |
| CMV, positive, n (%) | 8, (61%) | 9, (69%) | ns[#] |
| Age years, median [IQR] | 8 [3-12] | 42 [29-46] | p < 0.0001^ |

### Chai square
^ Mann-Whitney U test

#### Age is an intrinsic driver of inflammatory responses to malaria

**Supplementary Table S4: Antibody panel for ex vivo phenotyping**

| Antigen | Fluorochrome | Clone | Manufacturer | Cat # |
|---|---|---|---|---|
| CXCR3 | BV421 | 1C6 | BD Biosciences | 562558 |
| CD86 | BV480 | 2331 | BD Biosciences | 566131 |
| CD14 | BV510 | M5E2 | Biolegend | 301842 |
| CD127 | BV570 | A019D5 | Biolegend | 351308 |
| CCR6 | BV650 | 11A9 | BD Biosciences | 563922 |
| CXCR5 | BV711 | J252D4 | Biolegend | 356934 |
| HLA-DR | BV785 | L243 | Biolegend | 307642 |
| CD45RA | BB515 | HI100 | BD Biosciences | 564552 |
| CD3 | FITC | SK7 | Biolegend | 344804 |
| CD4 | PerCPCy5.5 | OKT4 | Biolegend | 317428 |
| CD19 | PE | HIB19 | Biolegend | 302208 |
| CD56 | PE-dazzle | HCD56 | Biolegend | 318347 |
| PD-1 | PE-CY7 | EH12.1 | BD Biosciences | 561272 |
| Vd2 | APC | B6 | Biolegend | 331418 |
| FoxP3 | AF647 | 206D | Biolegend | 320114 |
| CD16 | AF700 | 3G8 | Biolegend | 302026 |
| ICOS | APC-Cy7 | C398.4A | Biolegend | 301820 |
| Viability | NIR | | Invitrogen | L34975 |

**Supplementary Table S5: Antibody panel for innate cell parasite stimulation**

| Antigen | Fluorochrome | Clone | Manufacturer | Cat # |
|---|---|---|---|---|
| IL-12 | BV421 | C8.6 | BD Biosciences | 565023 |
| CD86 | BV480 | 2331 | BD Biosciences | 566131 |
| HLADR | BV570 | L243 | Biolegend | 307637 |
| IFNg | BV605 | B27 | BD Biosciences | 562974 |
| CD27 | BV650 | O325 | Biolegend | 302827 |
| CD45RA | BV711 | H100 | Biolegend | 304137 |
| TNF | BV750 | Mab11 | BD Biosciences | 566359 |
| CD3 | AF352 | UCHT1 | Invitrogen | 58003842 |
| IL-1b | FITC | CRM56 | Invitrogen | 11-7018-42 |
| CD14 | PerCPCy5.5 | M5E2 | Biolegend | 301823 |
| IL-10 | PE | JES3-9D7 | BD Biosciences | 559337 |
| CD56 | PE-dazzle | HCD56 | Biolegend | 318347 |
| IL-6 | PE-Cy7 | MQ2-13A5 | Biolegend | 501119 |

| | | | | |
|---|---|---|---|---|
| Granzyme B | APC | QA16A02 | Biolegend | 372203 |
| MCP-1 | AF647 | 5D3-F7 | BD Biosciences | 563496 |
| CD16 | AF700 | 3G8 | Biolegend | 302026 |
| Vd2 | APC-FIRE | B6 | Biolegend | 331419 |
| Viabilty | NIR | | Invitrogen | L34975 |

**Supplementary Table S6: Antibody panel for CD4 T cell parasite stimulation**

| Antigen | Fluorochrome | Clone | Manufacturer | Cat # |
|---|---|---|---|---|
| FOXP3 | BV421 | 206D | Biolegend | 320124 |
| CXCR3 | PAC Blue | G025H7 | Biolegend | 353723 |
| CD127 | BV570 | A019D5 | Biolegend | 351308 |
| CCR4 | BV605 | L291H4 | Biolegend | 359417 |
| CCR6 | BV650 | 11A9 | BD Biosciences | 563922 |
| CXCR5 | BV711 | J252D4 | Biolegend | 356934 |
| CCR7 | BV786 | 3D12 | BD Biosciences | 563710 |
| CD45RA | BB515 | HI100 | BD Biosciences | 564552 |
| Ki67 | FITC | B56 | BD Biosciences | 556026 |
| CD3 | AF352 | UCHT1 | Invitrogen | 58003842 |
| CD4 | PerCPCy5.5 | OKT4 | Biolegend | 317428 |
| PD-1 | PE-Cy7 | EH12.1 | BD Biosciences | 561272 |
| TNFR2 | AF647 | hTNFR-M1 | BD Biosciences | 562909 |
| CD25 | AF700 | 2A3 | BD Biosciences | 565106 |
| ICOS | APC-Cy7 | C398.4A | Biolegend | 301820 |
| Viabilty | NIR | | Invitrogen | L34975 |

### Age is an intrinsic driver of inflammatory responses to malaria

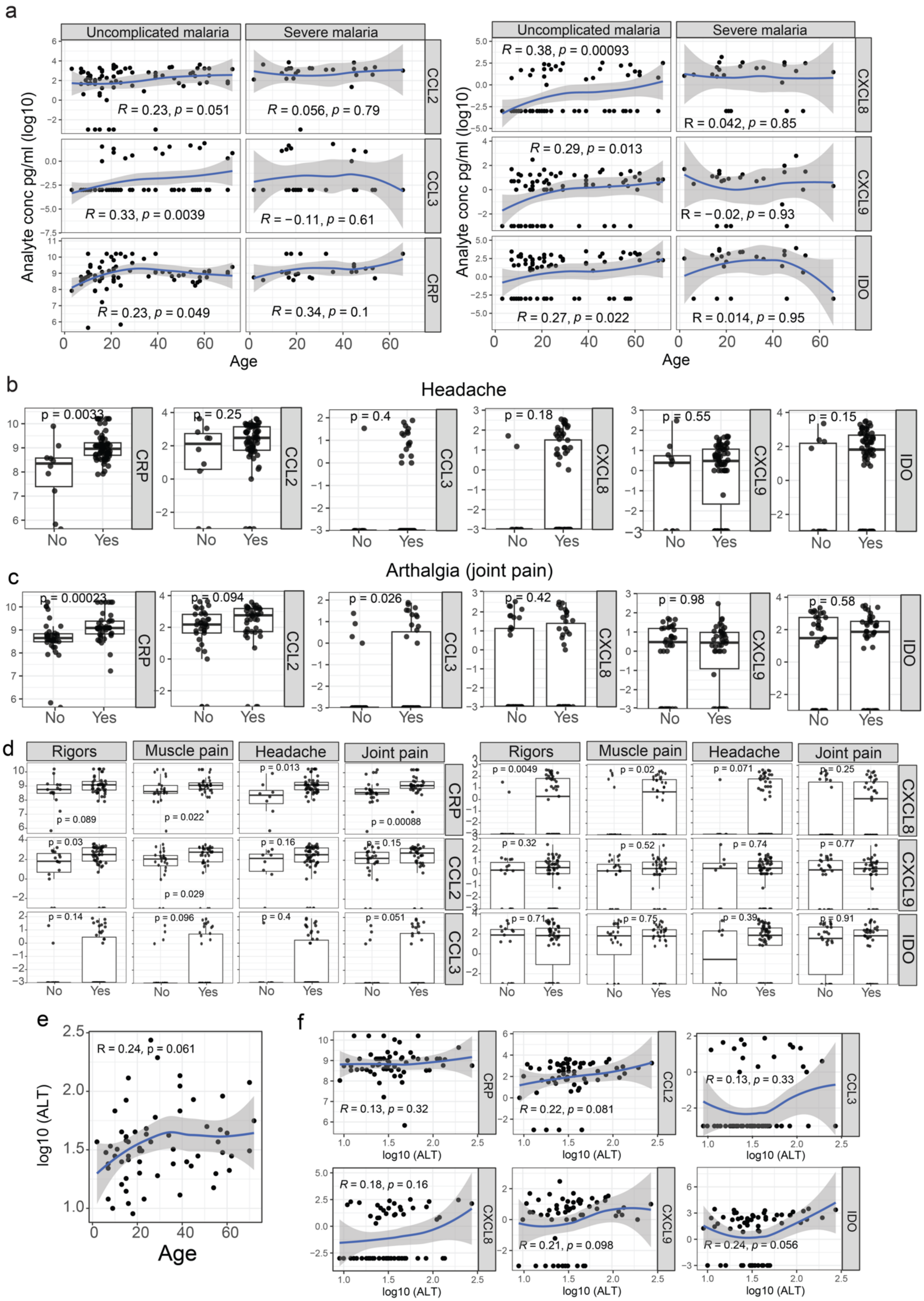

#### Age is an intrinsic driver of inflammatory responses to malaria

***Supplementary Figure 1. Associations between age, cytokines and symptoms.*** Thirteen analytes were measured in plasma from 97 individuals with acute malaria (male 78 (80%), age 21 [15-45] median [IQR])). **(a)** Age-associated inflammatory analytes stratified by malaria severity. Analyte concentration for patients with no symptoms (No) or with symptoms (Yes) **(b)** headache or **(c)** arthralgia (joint pain). **(d)** Analyte concentration for patients with no symptoms (No) or with symptoms (Yes) only in adults >12 years of age. **(e)** Correlation plot between ALT and age. **(f)** Individual correlation plots of analytes with ALT. Spearman's Rho and p are indicated. Tukey boxplots show the median, 25$^{th}$ and 75$^{th}$ percentiles. The upper and lower hinges extend to the largest and smallest values, respectively but no further than 1.5* IQR from the hinge. Dots represent individuals. Mann-Whitney U test was used for comparisons between no symptoms and yes symptoms.

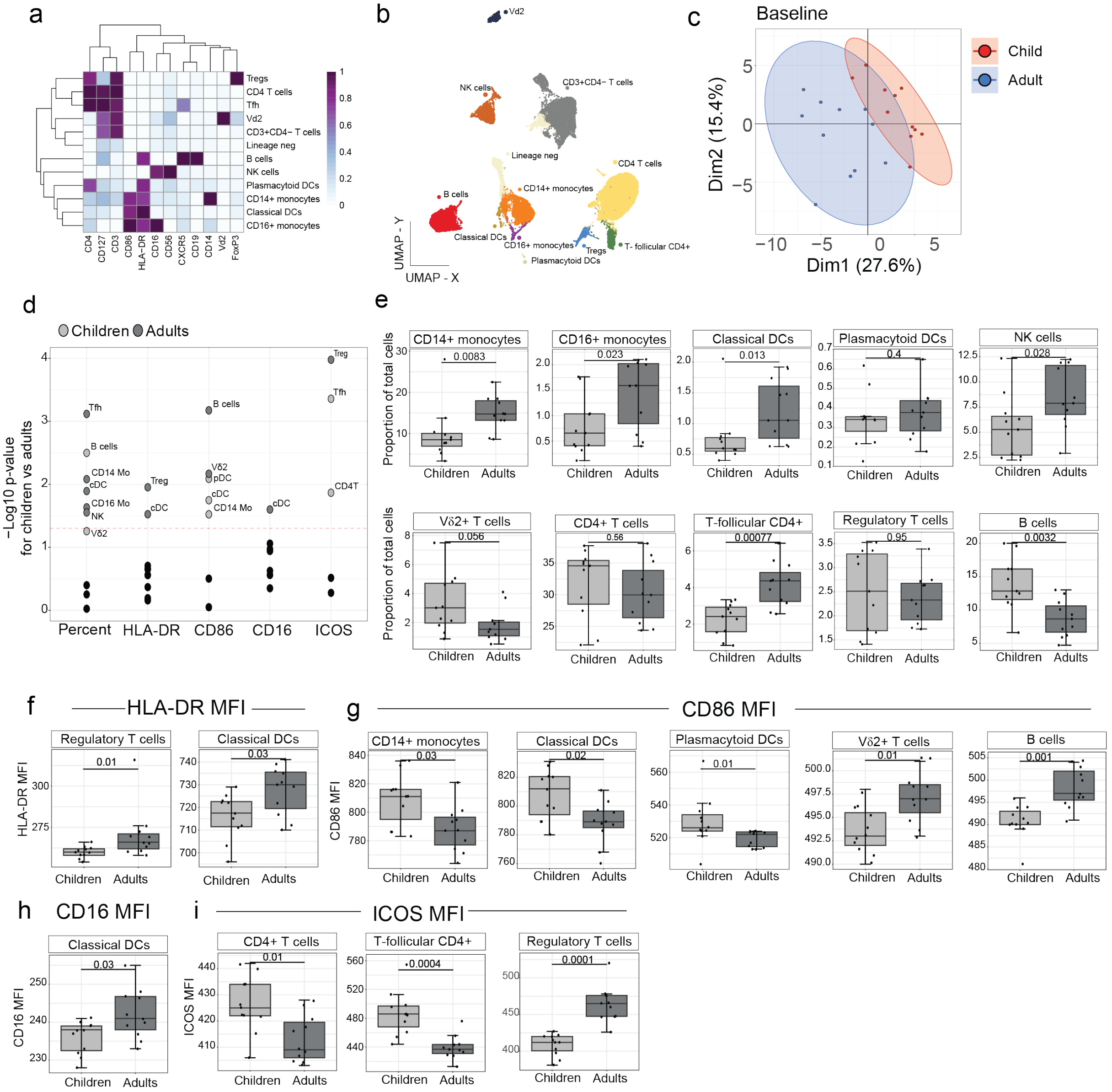


***Supplementary Figure 2. Frequency and phenotypes of malaria responsive immune populations in naïve children and adults.*** High-level cell lineages were identified by cell surface staining of malaria naive adult and children PBMCs using spectral flow cytometry. **(a)** Heatmap of median fluorescence expression (MFI) of protein markers within annotated PBMCs. Heatmap colour is normalised marker expression. **(b)** Uniform manifold approximation and projection (UMAP) of identified immune cell subsets in children (*n*=11) and adults (*n*=11). 11 distinct cell clusters (excluding lineage negative) were identified. **(c)** PCA of cell subset proportions and activation marker expression between children and adults. **(d)** Comparison of age specific differences between children and adults. The –log10 p-value calculated from Mann-Whitney test is indicated for the proportion (% of live PBMCs), HLA-DR MFI, CD86 MFI, CD16 MFI and ICOS MFI. **(e)** Immune cell type frequencies (as % of total cells), and significantly different **(f)** HLA-DR MFI, **(g)** CD86 MFI, **(h)** CD16 MFI, and **(i)** ICOS between children and adults. Tukey boxplots show the median, 25$^{th}$ and 75$^{th}$ percentiles. The upper and lower hinges extend to the largest and smallest values, respectively but no further than 1.5* IQR from the hinge. P values are Mann-Whitney U test.

#### Age is an intrinsic driver of inflammatory responses to malaria

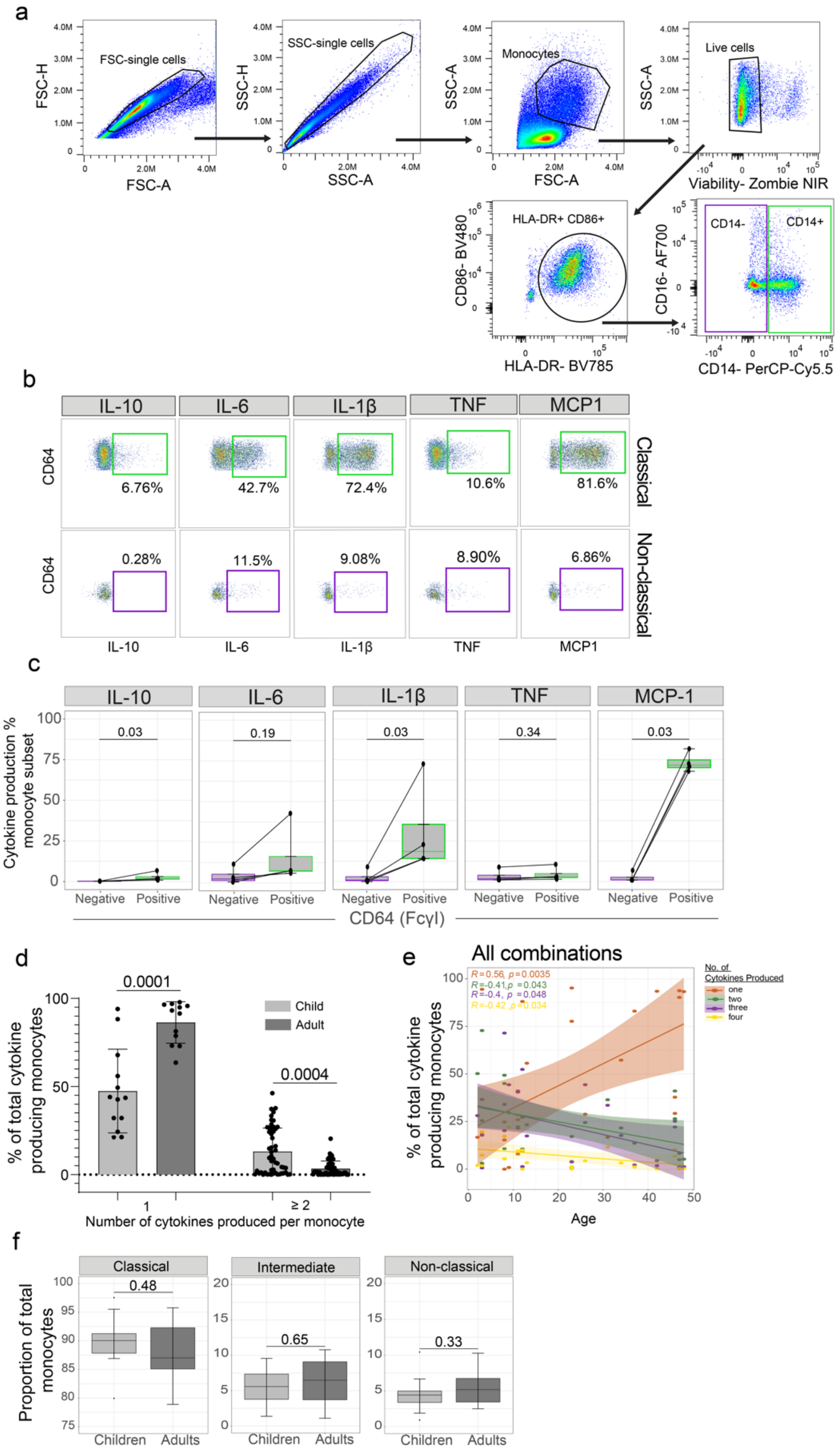

***Supplementary Figure 3*. *Functional cytokine response of classical monocytes.***
(**a**) Flow cytometry gating example for identifying classical and non-classical monocytes after stimulation. Total monocytes were identified as $HLADR^{+}$ and $CD86^{+}$, classical monocytes were identified as $CD14^{+}$ and non-classical monocytes were identified as CD14 -. To confirm the different functional potential of our identified monocyte subsets, we analysed their cytokine production in healthy adults (**b**) example flow cytometry plots (**c**) quantification of cytokine response from classical (green) and non-classical monocytes (purple). **(d)** Proportion of cytokine producing classical monocytes producing 1 or ≥ 2 cytokines. **(e)** Correlation between number of cytokines produced by monocytes and age. One cytokine is orange, two cytokines are green, three cytokines are purple, and four cytokines are yellow. Spearman's Rho and p are indicated. **(f)** Proportion of classical, intermediate and non-classical monocytes in children and adults. Tukey boxplots show the median, $25^{th}$ and $75^{th}$ percentiles. The upper and lower hinges extend to the largest and smallest values, respectively but no further than 1.5* IQR from the hinge. Lines represent paired observations, and negative and positive comparisons are Wilcox rank paired test.

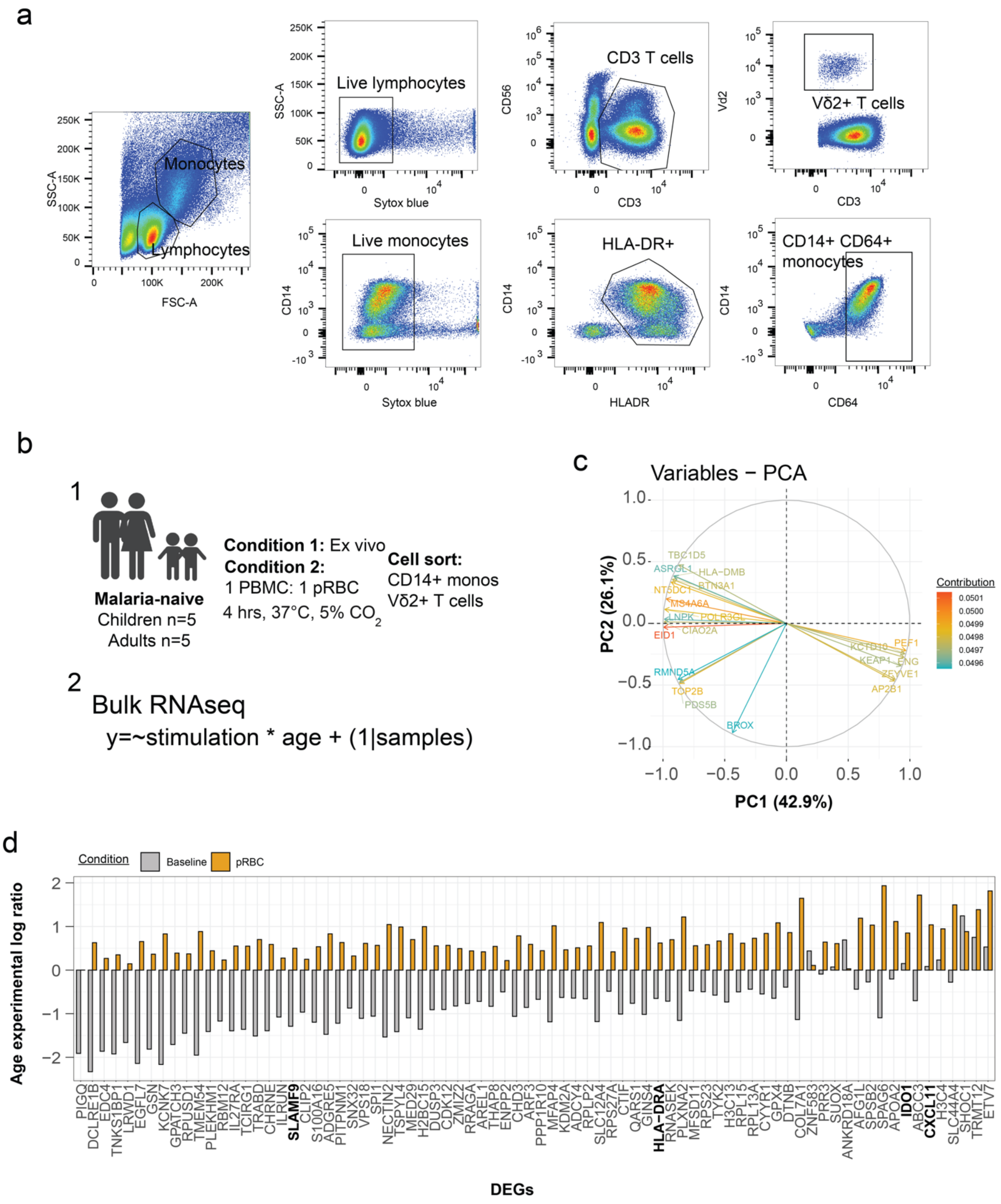

***Supplementary Figure 4. Classical monocyte bulk-RNA sequencing.*** **(a)**. Cell sort gating example, we identified classical monocytes as HLA-DR$^{+}$ CD14$^{+}$ and V$\delta$2$^{+}$ $\gamma\delta$ T cells as CD3$^{+}$ V$\delta$2$^{+}$. **(b)** Schematic of experimental (1) and analytical approach (2). CD14$^{+}$ classical monocytes were sorted ex vivo or following parasite stimulation from children (n=5) and adults (n=5). **(c)** Specific DEGs driving the variance for PCA1 and PCA2 in monocytes. **(d)** DEGs from classical monocytes with a qvalue (age & stim) < 0.001 (82 genes).

#### Age is an intrinsic driver of inflammatory responses to malaria

A positive age experimental log ratio for a DEG indicates it was higher in adults before or after stim and a negative age experimental log ratio for a DEG indicates it was higher in children before or after stim. Grey bars are unstim (baseline) values and orange bars are pRBC stim values. For example, CXCL11 (highlighted in blue) were higher in adults before and after stim (positive age experimental log ratio values).

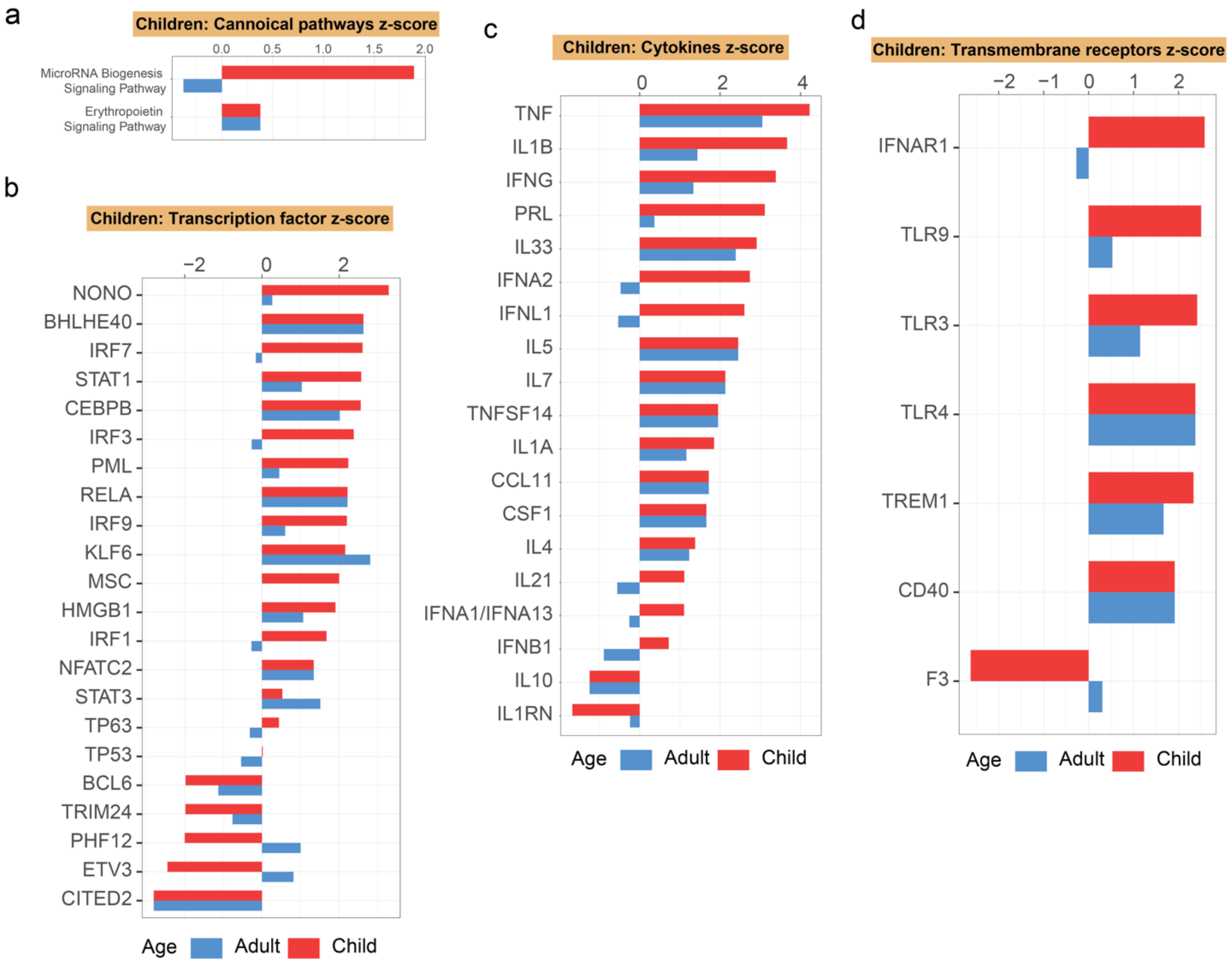


***Supplementary Figure 5. IPA analysis of monocyte genes which were higher in children compared to adults after primary pRBC stimulation.*** ***(*****a)** IPA performed using the log ratio value for children or adults and the FDR q-value of the DEGs that were upregulated following stimulation and were higher in children (red bars, figure 2 (d)). Significant pathways depicted. Predicted activated or inhibited upstream regulators analysis in adults and children **(d)** transcription factors, **(e)** cytokines, **(f)** transmembrane receptors, using DEGs as in a. Benjamin-Hochberg corrected P-values used to identify significant pathways and upstream regulators in the IPA analysis.

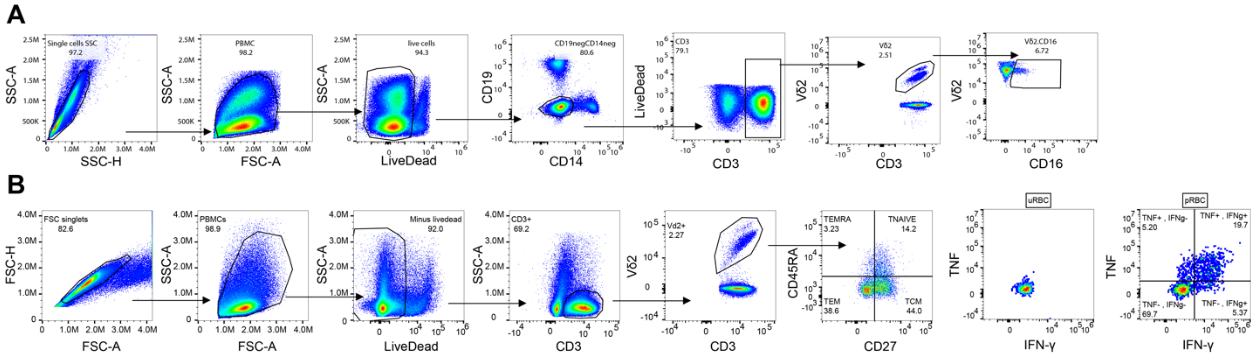


***Supplementary Figure 6. Representative gating strategy of V$\delta$2$^+$ $\gamma\delta$ T*** **cell** (**a**) *ex vivo* and after (**b**) *in vitro* stimulation. $\gamma\delta$ T cell memory subset defined by expression of CD27 and CD45RA (NAIVE: CD27$^+$ CD45RA$^+$, CM: CD27$^+$ CD45RA$^-$, EM: CD27$^-$ CD45RA$^-$ and EMRA: CD27$^-$ CD45RA$^+$). Single- and co-producing IFN$\gamma$ and TNF V$\delta$2 gates after uRBC and pRBC stimulation.

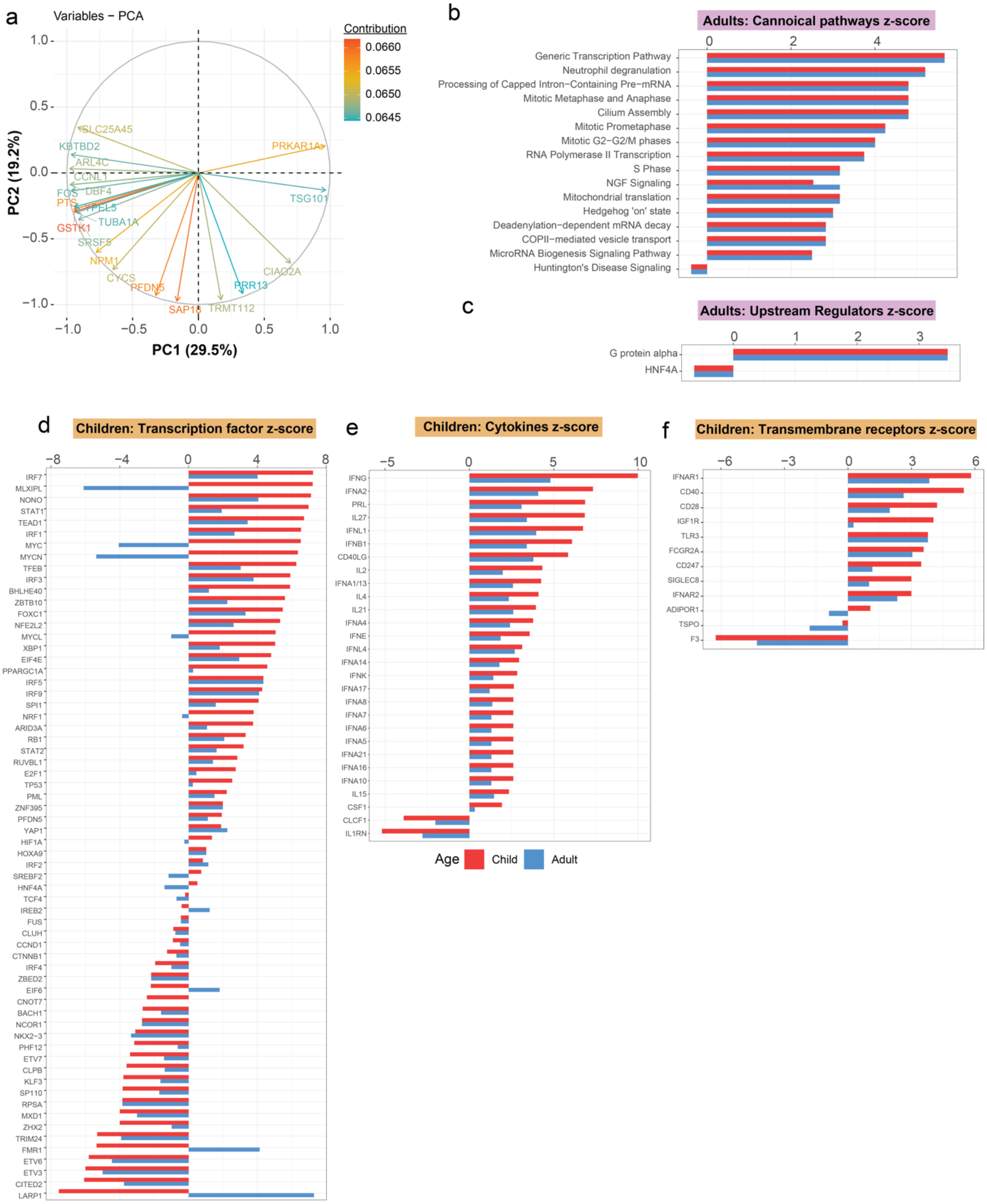


***Supplementary Figure 7. IPA analysis of Vδ2 T cell genes which were higher in adults compared to children after primary pRBC stimulation.*** **(a)** Specific DEGs driving the variance for PCA1 and PCA2 in Vδ2 T cells. **(b)** IPA performed using the log ratio value for children or adults and the FDR q-value of the DEGs that were upregulated following stimulation and were higher in adults (blue bars, figure 4 (d)).

Significant pathways depicted. **(c)** Predicted activated or inhibited upstream regulators analysis in adults and children, using DEGs as in a. Benjamin-Hochberg corrected P-values used to identify significant pathways and upstream regulators in the IPA analysis. Upstream regulator analysis for genes which were higher in children compared to adults after pRBC stimulation (orange bars) showing predicted **(d)** transcription factors, **(e)** cytokines and **(f)** transmembrane receptors to be activated or inhibited in adults and children, using DEGs as in Figure 4e.

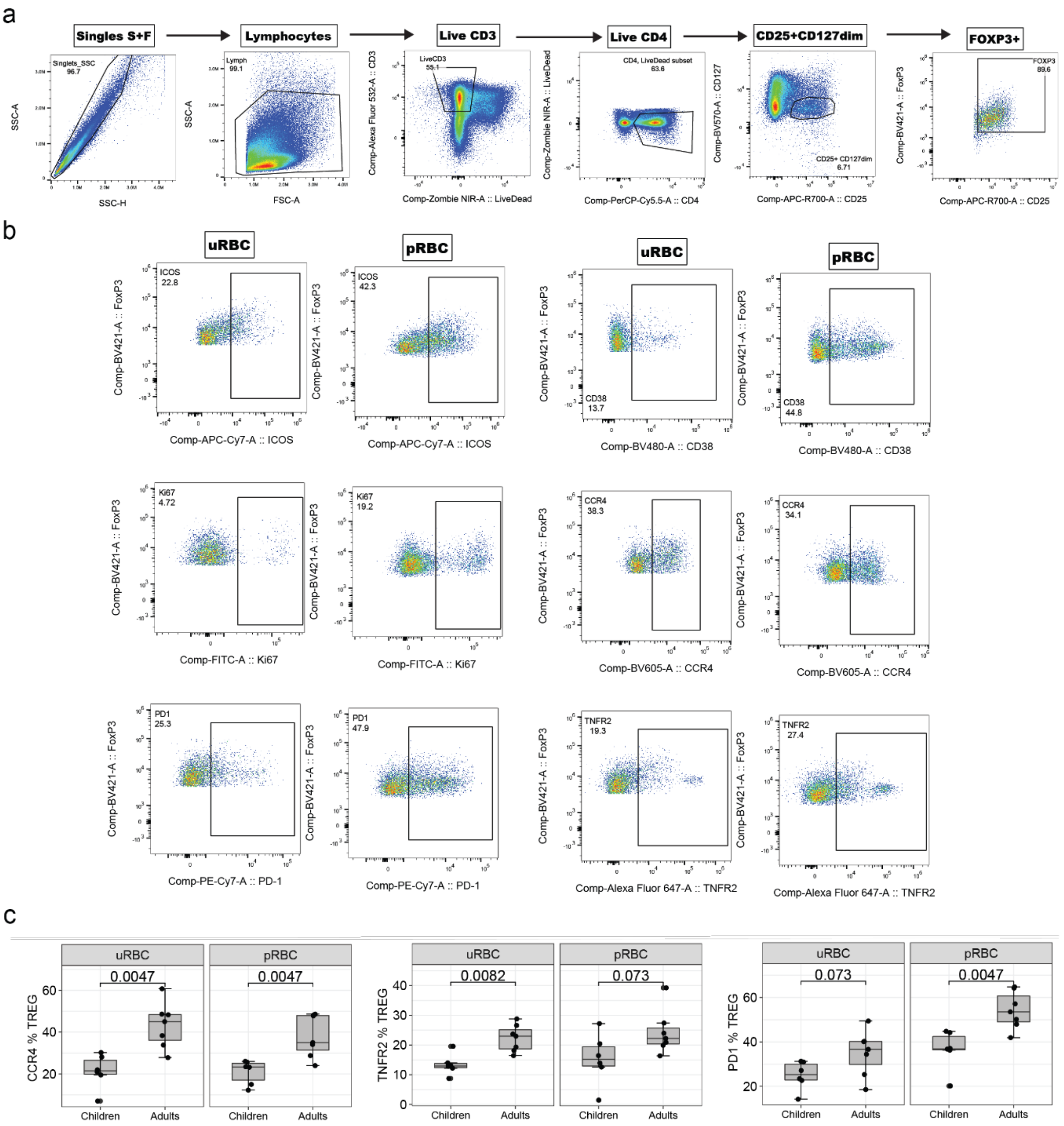


***Supplementary Figure 8. Identification of regulatory T cells in malaria naive children and adults.*** (**a**) Full flow cytometry gating strategy to identify CD25high CD127low FoxP3high Tregs. (**b**) Flow cytometry gating examples for activation and inhibitory marker expression on Tregs after uRBC and pRBC stimulation. (**c**) uRBC and pRBC age comparisons for inhibitory markers (CCR4, TNFR2, PD1).

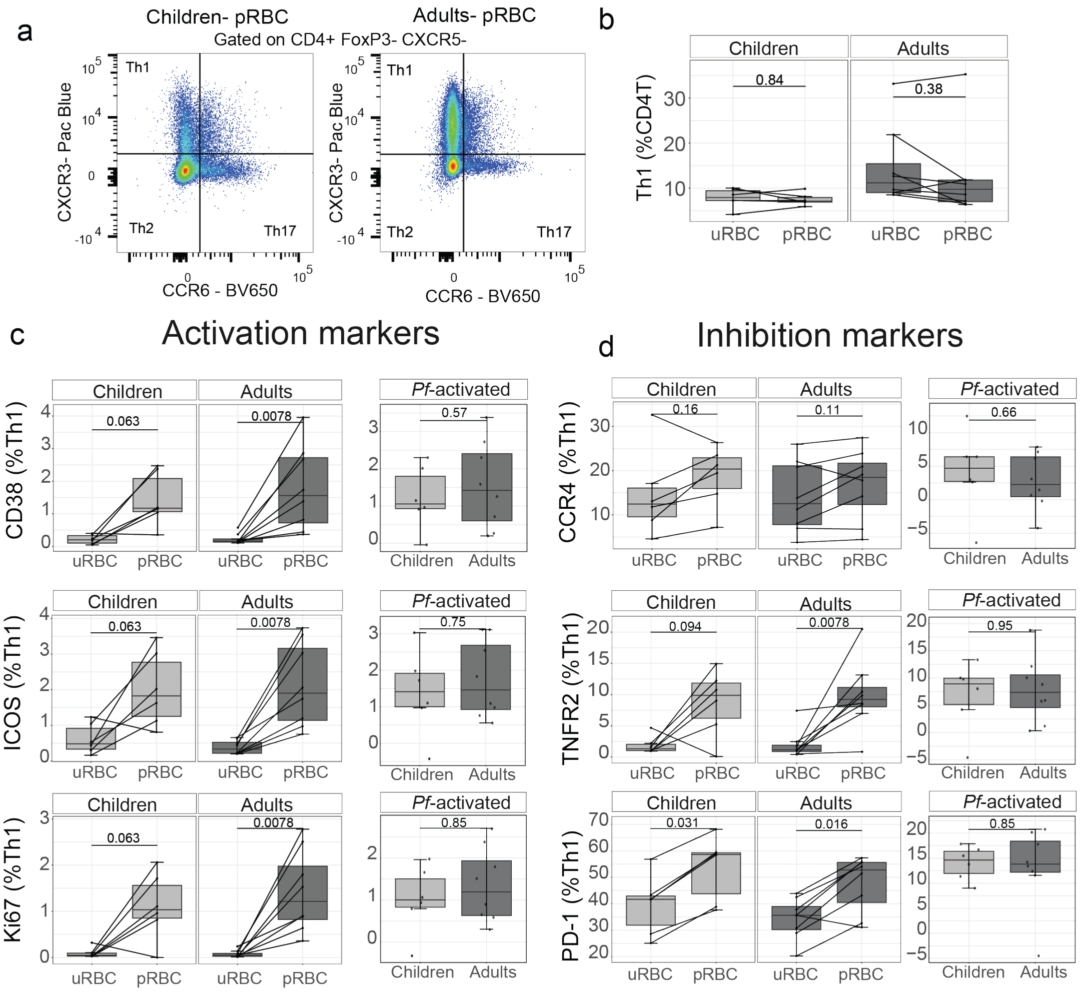


***Supplementary Figure 9. Comparable Th1 CD4$^+$ T cells response in both adults and children in response to malaria.*** Th1 CD4$^+$ T cell surface marker expression in children (*n*=6) and adults (*n*=7) was measured after 5 days *in vitro* co-culture with trophozoite stage *P. falciparum* red blood cells (pRBCs) and uninfected red blood cells (uRBCs). (**a**) Th1 CD4$^+$ T cells were identified as CXCR3$^+$ CCR6$^-$ CD4 T cells (FoxP3$^-$ CXCR5$^-$). **(b)** Th1 CD4$^+$ T cell frequency within CD4$^+$ T cells following stimulation. Surface marker frequency of inflammatory immune activation (**c**: CD38, ICOS and Ki67) and inhibition (**d**: CCR4, TNFR2 and PD-1) post stimulation of Th1 CD4$^+$ T cell frequency with pRBC, uRBC and *Pf* induced responses (quantified as expression in pRBC minus expression in uRBC cultured conditions). Tukey boxplots show the median, 25$^{th}$ and 75$^{th}$ percentiles. The upper and lower hinges extend to the largest and smallest values, respectively but no further than 1.5* IQR from the hinge. Lines represent paired observations. Wilcoxon signed rank test was used to compare paired data. Children and adult comparisons were made using the Mann-Whitney U test.
